# Golden Standard: A complete standard, portable, and interoperative MoClo tool for model and non-model bacterial hosts

**DOI:** 10.1101/2022.09.20.508659

**Authors:** Blas Blázquez, Jesús Torres-Bacete, David San Leon, Ryan Kniewel, Igor Martinez, Sandra Sordon, Aleksandra Wilczak, Sergio Salgado, Ewa Huszcza, Jarosław Popłoński, M. Auxiliadora Prieto, Juan Nogales

**Affiliations:** Department of Systems Biology, Centro Nacional de Biotecnología, CSIC, Madrid, Spain; Microbial and Plant Biotechnology Department, Biological Research Center-Margarita Salas, CSIC, Madrid, Spain; Interdisciplinary Platform for Sustainable Plastics towards a Circular Economy-Spanish National Research Council (SusPlast-CSIC), Madrid, Spain; Wrocław University of Environmental and Life Sciences, Department of Food Chemistry and Biocatalysis, Norwida 25, 50-375, Wrocław, Poland

**Author notes:** The authors wish it to be known that, in their opinion, the first five authors should be regarded as Joint First Authors.

## Abstract

Modular cloning assembly has become a benchmark technology in synthetic biology. However, there is a mismatch between its impressive development and the standardization required to promote interoperability between the different systems available. The full development of the field is thus hampered by a surge of oftentimes incompatible organism-specific systems. To overcome these issues, we present Golden Standard (GS), a Type IIS assembly method underpinned by the Standard European Vector Architecture (SEVA). GS unlocks modular cloning applications with any type of microorganism and delivers consistent combinatorial multi-part assembly of standardized genetic elements to create genetic circuits of up to twenty transcription units. Reliance on the Golden Gate syntax renders GS fully compatible with many existing tools and it sets the path towards efficient reusability of available part libraries and assembled TUs. GS was fully validated in terms of DNA assembly performance, portability and phenotype engineering in model and non-model bacteria. In order to facilitate the widespread adoption and future community-driven development of GS, we provide a web-portal featuring: i) a repository of parts and vectors, ii) a SBOLHub for exchange and analysis of constructs and iii) Wizard and Setup tools to guide the design of constructs using stored and user-specific parts.

## INTRODUCTION

Synthetic biology is a research field in continuous expansion and increased opportunities for new industrial processes (1). Synthetic biology aims at the construction of new genetic solutions to control cellular processes, which is achieved through rational design of transcription units (TUs) and, by extension, complex biological pathways and genetic circuits (2). The modular and hierarchical nature of biological designs reveals new possibilities for the development of rational and standardized mechanisms in order to improve the engineering process of specific biological solutions.

DNA assembly cloning is a widely used method to build synthetic genetic circuits. Traditionally, these cloning strategies were performed by digestion and ligation of DNA fragments (from genomic, plasmid or PCR origin), using BioBricks (3) or Gibson assembly (4). However, these approaches require specific designs for each step that can hamper complete standardization and parts reuse (3, 4). The Modular Cloning (MoClo) methodology has emerged as a powerful tool for standardizing the assembly of genetic parts. MoClo is based on Golden Gate cloning, which allows simultaneous and directional assembly of multiple DNA parts (5). This methodology was originally developed to work with plants, supporting fractionation of TUs into their basic parts (or level 0 parts): promoters, ribosome binding sites (RBS), coding sequences (CDS), terminators, etc. These modular elements can be further assembled into synthetic TUs (level 1 constructs), and more complex structures, such as genetic circuits (level 2 constructs) (6). To accomplish this, MoClo assembly combines Type IIS restriction enzymes and T4 DNA ligase in a one-pot reaction, thus making it possible to digest and assemble several DNA fragments simultaneously. Overall, Type IIS restriction enzymes are capable of cleaving any sequence outside of their recognition site, which opens up the possibility of designing compatible overhang ends (fusion sites) (7). Therefore, by placing the Type IIS enzyme recognition sites flanking level 0 parts it is possible to digest multiple DNA fragments while simultaneously assembling new DNA fragments in a directional, defined orientation to form level 1 TUs (8). The main DNA manipulation which is required when using MoClo technology is known as “domestication” of DNA parts and host vectors through the elimination of relevant Type IIS restriction enzyme sites. This simple process can be performed through PCR or by using chemically synthesized DNA parts.

The efficiency of MoClo technology as a cloning method for engineering genetic constructs has been demonstrated in a variety of diverse organisms, including bacteria (9–14), yeasts (15), plants (8, 16) and mammalian cells (17) (Table 1). In addition, the use of MoClo as a cloning system has seen an increase in the iGEM competition (13). Most of the MoClo systems use Type IIS enzymes that generate 4-nucleotide overhang fusion sites. Based on the fusion sites for level 0 parts described by Engler (8), BsaI has been established as the reference restriction enzyme for level 0. This common syntax and structure facilitates the construction of reusable and interchangeable part libraries and the development of standard protocols (18). This standardization effort has not prevented the appearance of alternative fusion site designs, thus increasing the variability of MoClo solutions. However, this increased scope has precluded full compatibility of the available libraries of parts and TUs categorized as level 0 and level 1, respectively. This incompatibility has also been augmented in some cases by the introduction of Type IIS enzymes that generate 3-nt overhangs (such as SapI) in the DNA parts, during the construction of TUs (11), and synthetic circuits (16).

**Table 1.** Modular cloning systems. Abbreviations: Prom, promoter; RBS, ribosome binding site; N-tag, N-terminal tag; CDS, coding sequence; C-tag, C-terminal tag; Ter, terminator. ^1^N. D: Not defined. ^2^N. A: Not applied.

| Name | Organism | Parts compatible with GS | Max. Parts assembled in TUs | Max. TUs | Ref |
| --- | --- | --- | --- | --- | --- |
| MoClo/ Golden Gate | Plants | Yes | **8** (Prom, RBS, N-tag x2, CDS, C-tag x2, Ter) | 17 | (6, 8, 19) |
| GoldenBraid | Plants, Fungi, Animalia | Yes | **11** (Prom, RBS, N-tag, CDS, C-tag, linker x5, Ter) | N.D.<sup>1</sup> | (20, 21) |
| CIDAR | *E. coli* | Yes | **4** (Prom, RBS, CDS, Ter) | 4 | (9) |
| EcoFlex | *E. coli* | No | **5** (Prom, RBS, N-tag, CDS, Ter) | 20 | (10) |
| Mobius assembly | *E. coli* | Yes | **10** (Prom, RBS, N-tag, CDS, C-tag, linker x4, Ter) | N.D.<sup>1</sup> | (22) |
| Midas | Fungi | Yes | **3** (Prom, CDS, Ter) | N.D.<sup>1</sup> | (15) |
| Start-Stop assembly | *E. coli* | No | **4** (Prom, RBS, CDS, Ter) | 15 | (11) |
| CyanoGate | Cyanobacteria | Yes | **7** (UpNeutral sites x2, Prom, CDS, Ter, downNeutral sites x2) | 4 | (12) |
| Multi-Kingdom | Fungi, Eubacteria, Protista, Plants, Animalia | No | **6** (Prom, RBS, N-tag, CDS, Ter, linker) | 5 | (23) |
| uLoop | Diatoms, yeast, plants, bacteria | Yes | **4** (Prom, RBS, CDS, Ter) | N.D.<sup>1</sup> | (24) |
| JUMP | Bacteria | Yes | **6** (Prom, RBS, N-tag, CDS, C-tag, Ter) | N.D.<sup>1</sup> | (25) |
| The Marburg Collection | Bacteria | Yes | 6 (Prom, RBS, N-tag, CDS, C-tag, Ter) | 25 | (13) |
| SEVAtile | Bacteria | No | 6 (Prom, RBS, CDS, Ter, RBS, Ter) | 1 | (14) |
| Golden Standard | Bacteria | N.A.<sup>2</sup> | 8 (Prom, RBS, N-tag x2, CDS, C-tag x2, Ter) | 20 | This work |

Despite most of the available MoClo systems having been specifically designed to engineer single species, new systems targeting more than one cellular host have recently been developed. In this sense, it is worth highlighting the Multi-Kingdom, JUMP and SEVAtile assembly systems (Table 1) (14, 23, 25). Multi-Kingdom proposes a universal system that supports construction of synthetic expression systems for both prokaryotes and eukaryotes. However, despite the fact that it uses a digestion/assembly system based on the alternation of Type IIS BsaI and BpiI enzymes, it is designed to use a unique syntax for fusion sites, thus reducing its operational standardization by limiting the possibility to directly exchange parts between different MoClo systems. JUMP assembly introduces a MoClo system for prokaryotic cells based partially on the architecture of SEVA vectors. This system uses the standard MoClo syntax to design fusion sites, but it relies on the alternative enzyme BsmBI instead of BpiI for level 2 assembly, which limits the interoperability among available MoClo systems. SEVAtile assembly also proposes a MoClo system based on the architecture of SEVA vectors. However, besides only being able to construct a single transcription unit with no possibility of including tags, it is limited by the impossibility to share parts due to the unique syntax used by this system.

Overall, a recurrent limitation and unsolved bottleneck in available MoClo systems is the absence of a meaningful community-guided development effort supporting individual users’ contributions to expand the universe of available DNA parts, receptor vectors or even novel applications. Furthermore, there is also a lack of intuitive computational frameworks to implement assembly of target TUs while providing the necessary rational, standardized and quantitative support to build complex metabolic pathways. Plant and cyanobacteria-based systems such as GoldenBraid (20) and CyanoGate (12) were designed with an eye on partially fulfilling these goals. However, they are clade or phyla-specific systems. Further efforts are needed to develop similar systems that are able to cover a broad range of organisms.

Here we present Golden Standard (GS), a standardized modular cloning technology that unifies and harmonizes available methods by extending the MoClo approach to a wide range of bacteria. This allows portability of the resulting final vectors for use in diverse microorganisms. Golden Standard adheres to SEVA standards, allowing not only modularity in the assembly process, but also modularization of the host vectors, thus resulting in a significant scope expansion. In order to facilitate widespread adoption and future community-driven development of GS, a web portal featuring the GS database, protocols and tutorials is available to make GS accessible to non-experts in modular cloning (http://sysbiol.cnb.csic.es/GoldenStandard/home.php).

## MATERIALS AND METHODS

Additional materials and methods are available in the Supplementary Information SI pdf file.

### Bacterial strains and plasmid construction

*Escherichia coli* DH5α and DH10B were used for all DNA assembly, unless otherwise stated. *E. coli* and *P. putida* were routinely grown on LB agar or in LB broth with shaking at 170 rpm at 37 °C and 30 °C, respectively. LB was supplemented with ampicillin (100 μg·mL^−1^), kanamycin (50 μg·mL^−1^), gentamicin (10 μg·mL^−1^) or streptomycin (75 μg·mL^−1^) as required. For PHB production in *E. coli*, LB was supplemented with glucose (10 g·L^−1^) and inoculated at OD_600_ 0.3 from an overnight culture of OD_600_~3. These cultures were grown for 24 hours at 37 °C and 200 rpm. Growth was monitored by measuring optical density at 600 nm (OD_600_) using a portable spectrophotometer (ThermoFisher Scientific).

The seven level 1 host vectors were constructed by standard cloning methods (Table S1). Briefly, MoClo fragments were amplified by PCR using primers with BpiI and BsaI sites and using the reporter LacZα as DNA template (Table S4). The PCR product was blunt ligated into pSEVA281, pSEVA231 and pSEVA221 plasmids digested with SmaI to divide the multicloning site into a prefix (PacI-SmaI) and a suffix (BamHI-SpeI). To remove a BpiI site from the pBBR1 origin of replication, pBBR1 was amplified in two fragments using the oligo pairs 5pBBR1_1 - 3pBBR1_1 and 5pBBR1_2 - 3pBBR1_2 that include overlapping regions with a single nucleotide mutation to eliminate the BpiI site. The two PCR products then served as a template for overlap extension PCR to generate a fragment containing the domesticated origin of replication. This fragment was used as pBBR1 origin of replication. Level 2 and level 3 host vectors were constructed following the same method used for the level 1 vectors (Table S1).

### Construction of level 0 parts

Fragments were amplified by PCR, oligonucleotide annealing or by chemical synthesis while introducing silent mutations (when appropriate) to eliminate restriction enzyme sites used in the assembly system. If subsequent traditional restriction enzyme cloning is planned, restriction sites present in the SEVA backbone were also removed. Further details can be found in the supplementary materials and methods.

### Modular assembly protocol

Assembly reactions were set up using 40 fmol of each donor plasmid part, 200 U T4 ligase (NEB), 10 U BsaI-HFv2 (NEB) for assembly into levels 1 and 3 or 10 U BpiI (Thermo Scientific) for assembly into levels 2 in 20 μl 1x T4 ligase buffer (NEB). The reaction condition was: 28 cycles of 37 °C 1.5 min plus 16 °C 3 min, then 50 °C 5 min, 37 °C 5 min, and 80 °C 10 min. Reactions were then transformed into *E. coli* DH5αor DH10B, and plated on LB agar with antibiotic plus 40 μg/ml X-Gal and incubated at 37 °C overnight. White colonies were selected and screened by DNA sequencing.

### Analysis of GFP reporter by flow cytometry

Level 1 GFP reporter plasmids were electroporated into *P. putida* KT2440 (52). Isolated clones of *E. coli* DH5α and *P. putida* KT2440 containing the level 1 GFP reporter were used to inoculate 10 mL LB broth supplemented with kanamycin. Overnight cultures were diluted 1:1000 into 10 ml fresh medium, and grown until the cultures reached OD_600nm_ 1.2. Cultures were diluted 1:50 in sterile PBS and immediately subjected to flow cytometry analysis.

GFP Fluorescence was measured using a MACSQuant™ VYB cytometer (Miltenyi Biotec, Bergisch Gladbach, Germany), using a λ_Excitation_ 488 nm and λ_Emission_ 525 nm. Two biological replicates were performed with the collection of 50000 events gated by forward scatter height (FSC-H) and side scatter height (SSC-H), the median from the detector was exported as the fluorescence value. Flow cytometry data were analysed using FlowJo (www.flowjo.com).

### Polyhydroxybutyrate (PHB) quantification and monomer composition

PHB monomer composition and PHB content were determined by Gas Chromatography-Mass Spectrometry (GC-MS) of the methanolysed polyester (53) (see supplementary material and methods for details).

### Analysis of expression of level 1 N- and C-tagged OleD glucosyltransferase

*E. coli* were electroporated with a series of level 1 OleD GT vectors with different N-terminal purification tags and/or C-terminal GFP (see supplementary information, Table S2). Isolated clones were used to inoculate 1 mL (mixed assembly) or 25 mL (individual assemblies) of LB broth supplemented with kanamycin and cultured overnight at 37 °C with shaking at 120 rpm. Cultures were subsequently harvested at 4000 xg for 30 min and cell pellets were resuspended in 0.5 mL (mix) or 5 mL (individual) of lysis buffer (30 mM Na phosphate pH 8.0, 300 μg/mL lysozyme), sonicated and centrifuged 20000 xg for 30 min at 4 °C. The resulting supernatant was directly used in further analysis of protein concentration (individual assemblies), fluorescence, glucosyltransferase activity, SDS-PAGE (individual assemblies) and protein purification (individual assemblies, N-His6 version only). Protein concentration was evaluated by Bradford assay using BSA as the reference for calibration (54). Fluorescence was measured using a Cary Eclipse Varian fluorimeter with unmodified OleD crude extract as a reference. Glucosyltransferase relative activity was evaluated via quick HPLC assay (for details of sample preparation, product and analytical procedure see supplementary information), using 100 μL of crude supernatants in a sodium phosphate buffer (pH 8.0, 30 mM) supplemented with 0.05 mM xanthohumol (10 μL of DMSO stock solution) as glucose acceptor and 0.1 mM UDP-Glucose as glucose donor in a final reaction volume of 1 mL for 20 min at 30 °C and 800 rpm using a Thermoshaker (Eppendorf). Crude extracts used to test individual assemblies were diluted to an exact protein concentration of 50 μg/mL. IMAC protein purification was performed on a GE Healthcare His SpinTrap column following the manufacturer’s instructions. SDS-PAGE was performed using standard conditions with Laemmli denaturing buffer, precast TGX gels (Bio-Rad) and Broad Range Protein Marker (New England Biolabs).

## RESULTS

### Golden Standard features, transcription units and circuit assemblies

GS is a flexible synthetic biology tool for modular cloning based on PhytoBricks parts (6). In addition, GS adopts the standardized vector structure provided by SEVA (26). As with other Golden Gate-based DNA assembly systems, GS is based on the hierarchical assembly of functional DNA parts into increasingly complex genetic circuits using receptor vector levels 0 to 3.

To build the basic level 0 parts (promoters, RBS, CDS, terminators, N-, and C-terminal tags), DNA fragments can be created by PCR amplification, oligonucleotide annealing, or by chemical synthesis. Irrespective of the method used to generate the DNA fragment, the sequence must be domesticated in order to eliminate BsaI and BpiI recognition sites. It is also strongly recommended to eliminate the restriction sites that define the different modules of the SEVA vectors (27) if further traditional cloning steps are envisioned. GS level 0 parts were cloned into the acceptor vector pSEVA182 linearized with SmaI to facilitate blue-white selection of positive clones. Furthermore, parts from other MoClo systems with the same fusion site syntaxes can be used directly as GS parts, including level 0 parts from MoClo/Golden Gate, CIDAR, Mobius or CyanoGate (6, 8, 9) (Table 1), see below.

GS level 0 parts are flanked by BsaI restriction sites with unique 4-nucleotide fusion sites, labelled A to I depending on their category (Figure 1). By concatenating compatible fusion sites, multiple parts can be assembled in order to create customized TUs (Figure 1). By using a one-pot restriction and ligation reaction, correctly assembled parts lose the BsaI site while re-ligated fragments are re-cleaved again in each new cycle. Once the level 0 parts are assembled into TUs, no BsaI sites remain in the final level 1 construct, thus achieving highly efficient assembly (≥95%, Figure S1). Basic functional GS level 1 TUs are made up of four level 0 parts: promoter, ribosomal binding site (RBS), coding sequence (CDS) and terminator, assembled within a level 1 host vector between fusion sites A to I, and using the combination of fusion sites A-B, B-D, D-G and G-I (Figure 1). To assemble more complex TUs requiring additional functional parts, such as signal peptides, anchors or tags, there are four additional positions available: two for N-terminal (C and E) and two for C-terminal (F and H) additions that can be used to deliver the final construct between the fusion sites A to I (Figure 1). Overall, up to eight level 0 parts can be combined and assembled into different level 1 receptor vectors.

**Figure 1.**
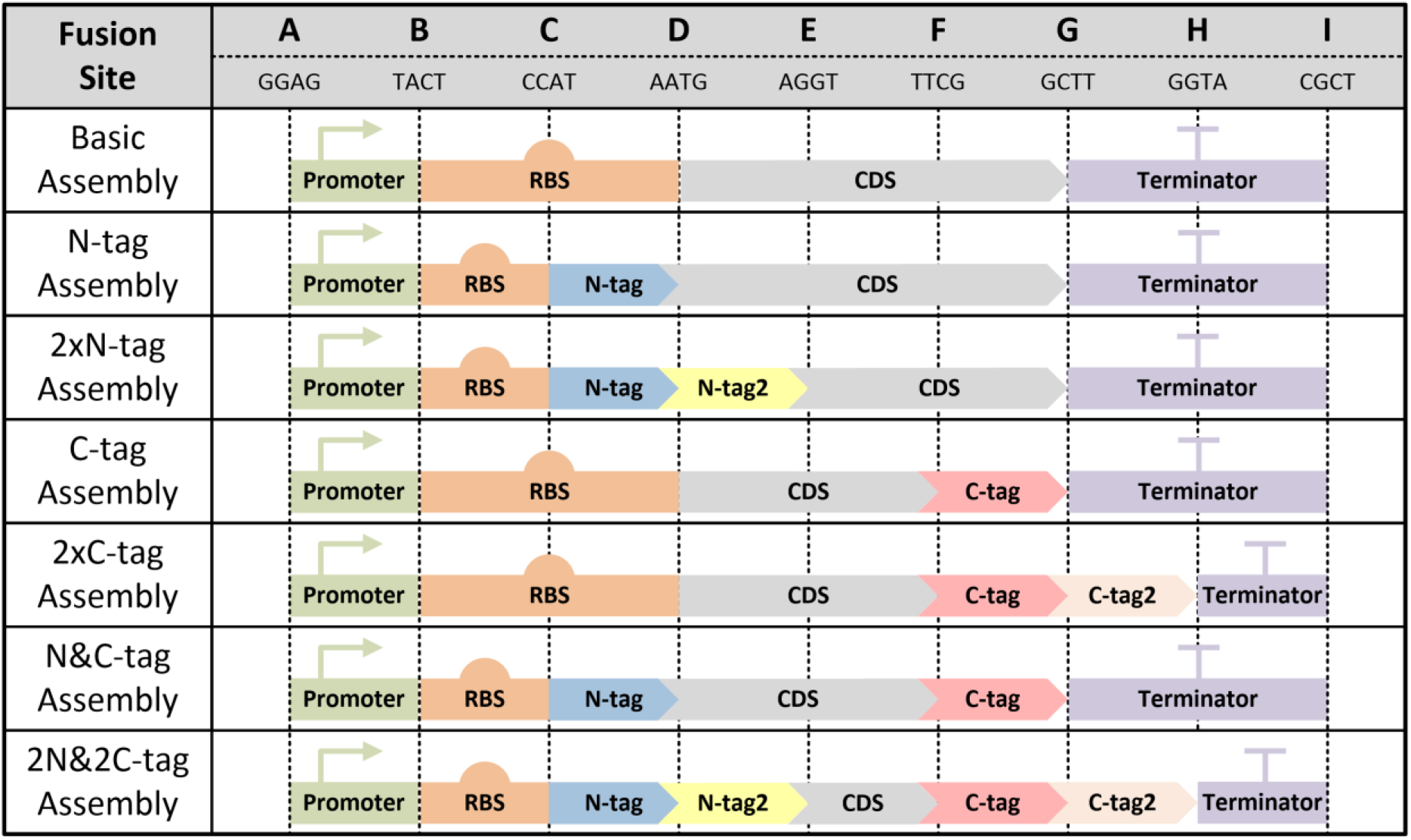
Golden Standard fusion sites. Modules can be combinatorially assembled into a level 1 host vectors to make a transcription unit.

Depending on the future position of the TUs in more complex circuits, there are six level 1 receptor vectors available (Figure 2). Additionally, we also created special level 1 host vectors with inverted I to A fusion sites to construct TUs with the opposite transcription orientation. We initially included three sets of level 1 host vectors covering a large range of origins of replication, RK2, pBBR1 and pUC for low, medium and high copy number plasmids, respectively (Table S1). The broad range of origins of replication (RK2 and pBBR1) provide compatibility with Gram-negative bacteria other than *E. coli*, such as *Pseudomonas*, *Halomonas*, *Acinetobacter*, *Bordetella, Rhizobium, Rhodospirillum, Xanthomonas*, etc., thus providing a significant advantage with respect to other MoClo systems (9–11, 22). Level 1 host vectors contain a kanamycin resistance cassette and the *lacZ* gene to facilitate the selection of colonies by alpha complementation. Flanking the BsaI sites in level 1 constructs there are additional type IIS BpiI sites to allow subsequent assembly of multiple TUs in level 2 host vectors (Figure 2, Figure 3).

**Figure 2.**
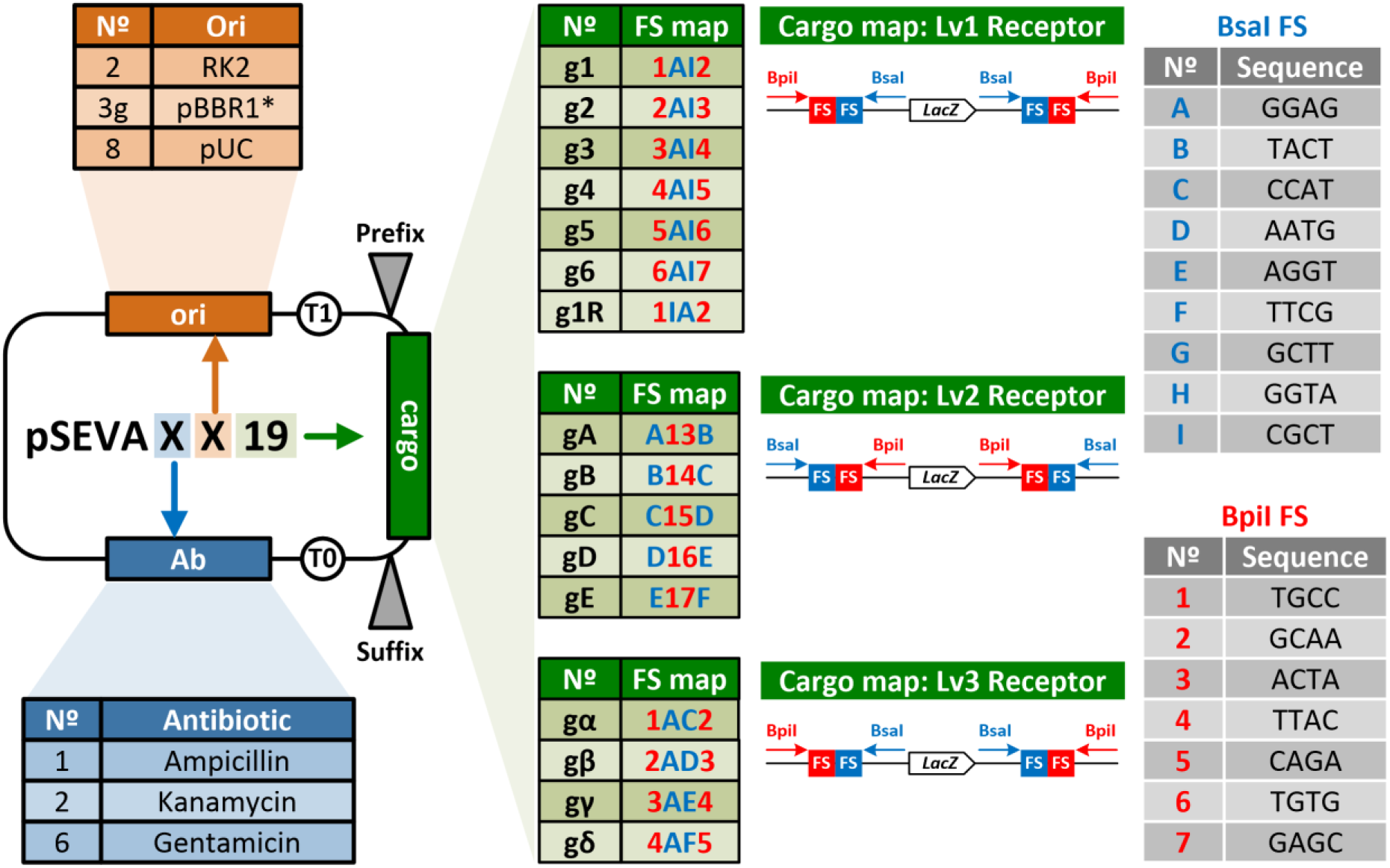
Structure and nomenclature of GS pSEVA vectors (Table S1). Following SEVA nomenclature, GS pSEVA vectors are named by 3 digits where the first number denotes the antibiotic resistance marker (blue), the second number identifies the origin of replication (orange), and the third number identifies the cargo (green). Golden Standard cargo is denoted with the number 19. In addition, vector level 1 (Lv1) receptors, level 2 (Lv2) receptors and level 3 (Lv3) receptor are named with the letter g followed of numbers, capital letters and Greek letters, respectively. The position of the fusion sites (FSs) of each specific vector and the basic structure of the cargos of each type of receptor vector (Lv1, Lv2 and Lv3) is indicated. Sequences of the BsaI FSs (blue) and BpiI FSs (red) are shown in the right tables.

Five level 2 vectors are available, thus supporting assembly of up to six TUs. The assembly of level 2 constructs is driven in a fixed and directional order through the fusion sites generated by BpiI digestion numbered 1 to 7 (Figure 2, Figure 3). The choice of the final level 2 host vector will depend on the number of transcription units to be assembled in the final construct. In contrast to other MoClo systems with a single level 2 receptor vector that require linker parts to fill empty positions, the library of level 2 vectors provided by GS significantly increases assembly efficiency by avoiding the use of extra adapters. The level 2 host vector collection includes plasmids with RK2, pBBR1 and pUC origins of replication and the gentamycin resistance cassette in order to facilitate selection (Table S1). These vectors also contain BsaI sites external to the BpiI sites in order to allow fusion of multiple transcription units into level 3 host vectors (Figure 2, Figure 3). We constructed four level 3 host vectors containing BsaI fusion sites to assemble between seven and twenty TUs using the appropriate linkers, all of them featuring the kanamycin resistance cassette (Figure 2).

**Figure 3.**
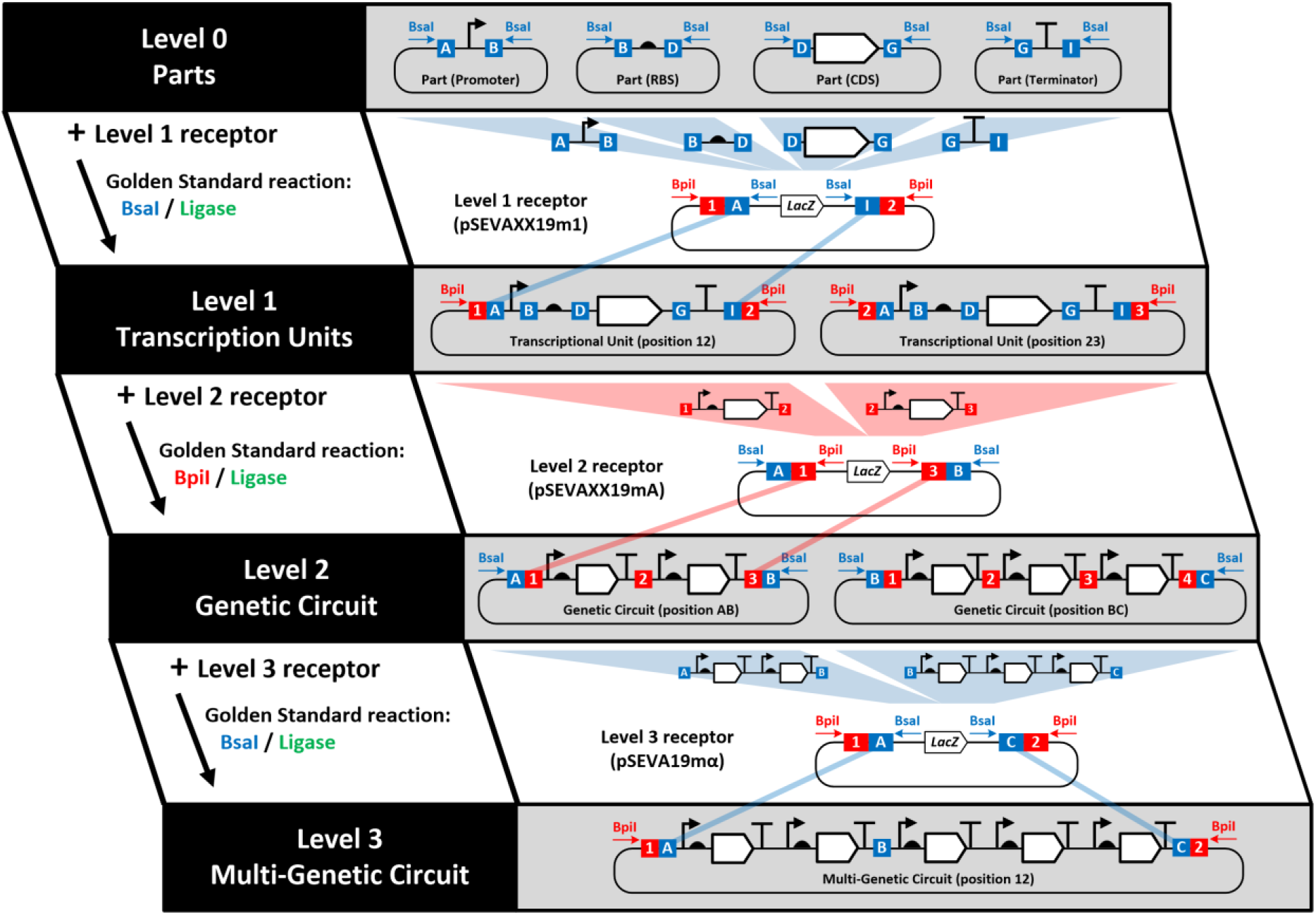
Hierarchy and graphical description of Golden Standard assembly. Hierarchy of Golden Standard assembly ranges from basic DNA parts (level 0), which are assembled to generate transcription units (level 1), which in turn can be assembled into more complex genetic circuits (level 2 and level 3). A Golden Standard reaction consists of a series of restriction/ligation cycles, whose operation is the same, regardless of the level under construction. Restriction steps release the genetic features flanked by the corresponding Type IIS enzyme (BsaI or BpiI), and removes the lacZ gene from the receptor vector. Ligation steps assemble the genetic features in a pre-defined order into the receptor vector.

In addition, all receptor vectors (levels 1 to 3) included the SEVA multicloning site divided into a prefix (PacI to SmaI sites) and suffix (BamHI to SpeI sites) flanking the MoClo cargo, thus allowing further pathway/circuit modifications by traditional restriction enzyme cloning (Figure 2). Furthermore, the adoption of SEVA standards allows the *ad hoc* construction of novel level 1-3 receptor vectors to extend GS to alternative non-model bacteria.

To increase the flexibility and reusability of GS, we additionally created linker DNA parts (short non-coding DNA sequences) that can be used to de-functionalize any level 0 part or level 1 TU. The linkers are available as level 0 (BsaI and their corresponding fusion sites) and level 1 (BpiI and their corresponding fusion sites). The inclusion of multiple linkers allows the construction of polycistronic circuits (by replacing promoters and/or terminators) and the generation of genetic circuits of up to twenty transcription units, among other applications (a detailed protocol describing the construction of these complex circuits is provided in the supplementary materials and methods, Figure S2).

GS is initially launched with a library of: i) seven constitutive and six inducible promoters; ii) three mono and two bicistronic RBSs; iii) six N-terminal and three C-terminal tags; iv) four widely used CDS such as fluorescent reporter genes; v) five terminators; vi) fifteen linkers for construction of TUs and assembly of level 2 and 3 circuits; vii) three preassembled transcription factors in order to allow the use of inducible promoters and viii) a collection of forty-one host vectors. The list of ninety-six available plasmids for distribution (including receptor plasmids and parts) can be found in Table S1.

### Expanding vector portability, bacterial hosts and gene expression control through Golden Standard assembly

Portability is the capability to exchange biological parts across organisms regardless of the origin of the DNA sequence. In the frame of synthetic biology and DNA assembly, portability is one of the key features allowing part reusability (28). Among other benefits, portability allows saving the vast effort required to generate and characterize parts for each individual organism. Despite the large effort to make highly portable libraries of DNA parts in the context of MoClo systems, libraries of receptor vectors suitable for use in a broad range of bacterial hosts are just starting to be developed (23, 29).

By adopting SEVA standards, GS supports the direct expansion of MoClo approaches to non-model organisms by avoiding dependence on shuttle vectors (30). To demonstrate the portability of the GS system, we assembled a TU expressing the reporter gene *gfp* under the control of the constitutive Pem7 promoter in receptor vectors with different origins of replication. This approach allowed for increased copy numbers (RK2, pBBR1 and pUC) (Figure 4A). We then tested the performance of these expression plasmids in both model (*E. coli*) and non-model (*P. putida*) microorganisms by monitoring GFP-based fluorescence using flow cytometry (Figure 4B & 4D). As expected, constructed plasmids could be used interchangeably in both biotechnology-relevant strains. The expression profiles were consistent with the expected number of copies provided by each origin of replication (31). In fact, we found a linear correlation between the plasmid copy number and GFP fluorescence returning a 0.9841 fit in *E. coli* (Figure 4C). High correlation (0.9931) was also found for complex phenotypes such as polyhydroxybutyrate (PHB) production (Figure 4C see below Figure 5). This approach make evident the host-dependent expression of GFP irrespective of the plasmid used and we found lower expression levels in *P. putida* (Figure 4B & 4D). These differences could be explained in terms of lower copy number of plasmids in *P. putida* and/or as a direct consequence of the biological context surrounding the genetic device, as recently shown (32). Therefore, adequate monitoring of the performance of a given part, TU, and/or circuit in the target bacterial chassis requires accurate measurements. Critically, GS largely facilitates this important task in synthetic biology.

**Figure 4.**
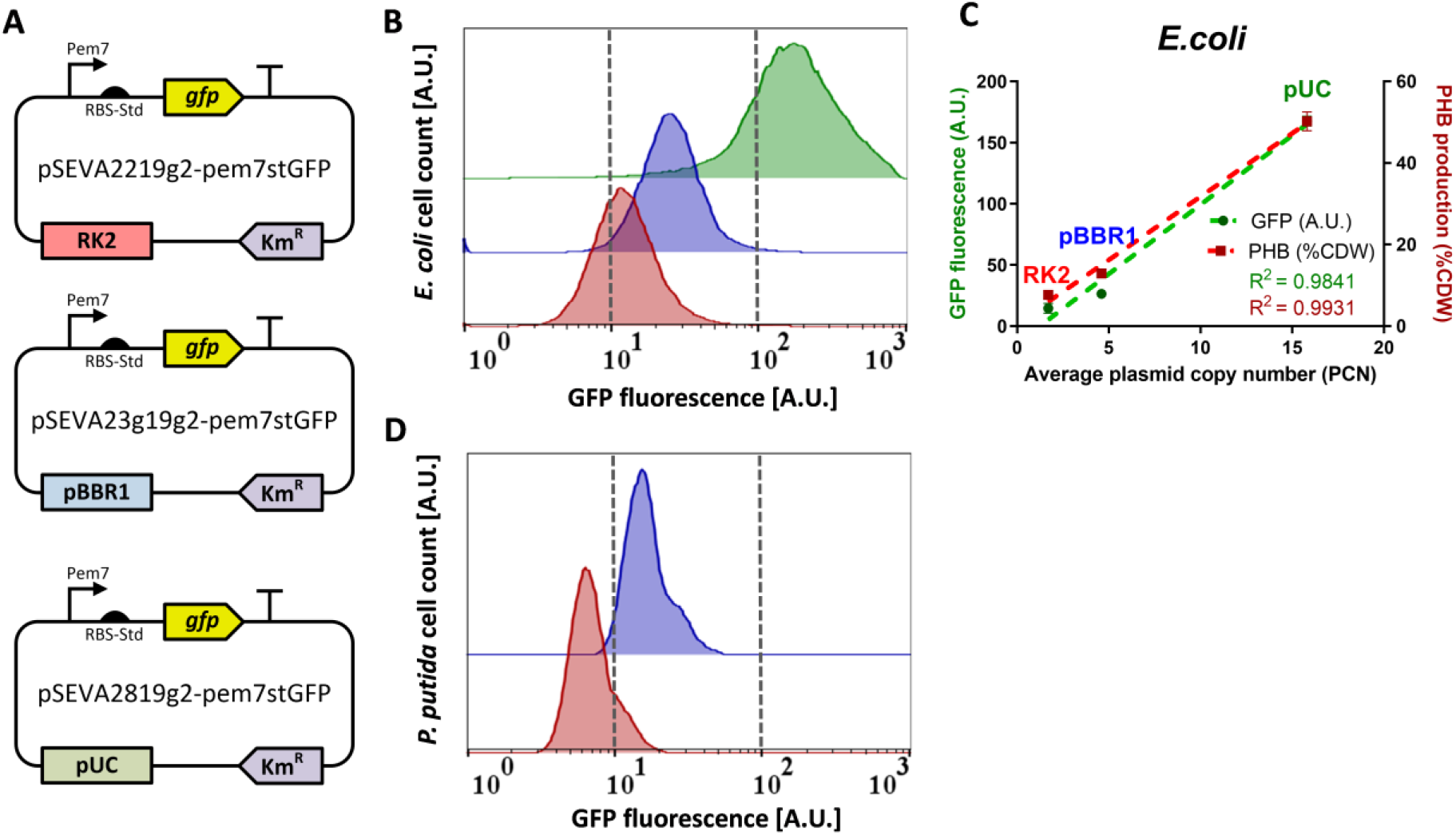
Portability of Golden Standard assembly. A) The gfp gene was assembled using the constitutive Pem7 promoter, a consensus RBS (RBS-St) and the iGEM BBa_B1006 terminator in level 1 vectors with different origins of replication. GFP fluorescence was analysed via flow cytometry. B) GFP intensity of E. coli cells carrying plasmids with RK2 (red), pBBR1 (blue) or pUC (green) origins of replication. C) GFP intensity and PHB production in E. coli cells is dependent on plasmid copy number. D) GFP intensity of P. putida cells carrying plasmid with RK2 (red) or pBBR1 (blue) origins of replication. For cytometry representation, representative samples were taken from a batch cultivation at OD_600_ of 1.2.

**Figure 5.**
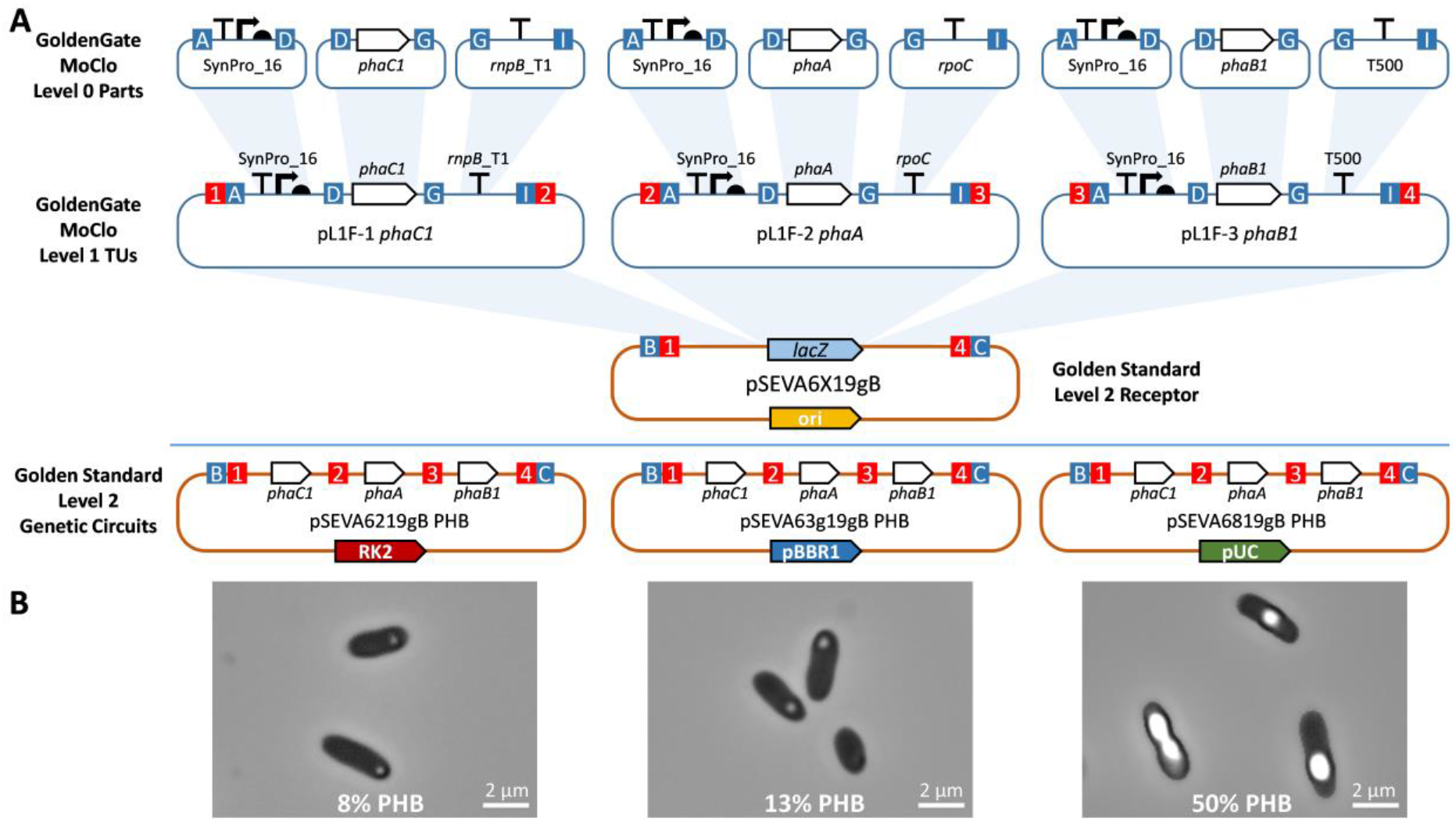
A) PHB operon construction combining Golden Standard with Golden Gate. Level 0 parts for PHB production previously constructed with the Golden Gate-MoClo toolkit were assembled in level 1 Golden Gate-MoClo vectors to obtain three TUs. Afterwards, these TUs were assembled in three level 2 GS pSEVA plasmids with RK2, pBBR1 or pUC origins of replication. B) PHB granules present in E. coli DH10B cells harbouring plasmids pGS6219gB PHB (8% PHB), pGS63g19gB PHB (13% PHB) and pGS6819gB PHB (50% PHB). Image captured using phase contrast microscopy.

On the other hand, the broad library of GS receptor vectors unlocks a new layer of protein expression optimization during the combinatorial assembly of a given metabolic pathway. Traditionally, optimal pathway expression has been addressed by tuning the strength of the genetic parts involved in transcription and translation (e.g. promoters, RBS, etc.) regardless of the number of copies of the plasmid in the cell host. This is because most available Golden Gate systems are based on vectors with a unique origin of replication, mainly pUC. This feature excludes plasmid copy number as an effective variable that could be tuned for gene expression optimization. With GS, this additional layer of gene expression optimization becomes available and delivers a large advantage over previous Golden Gate systems because it supports the construction of plasmid libraries displaying broad ranges of expression performance by using vectors from across a range of copy numbers to assemble part libraries.

### Golden Standard provides high interoperability with available MoClo systems

In addition to portability, interoperability and compatibility with existing platforms becomes an essential feature of any genetic device when it comes to facilitating community-driven development of increasingly complete platforms, while at the same time promoting reusability of parts and plasmids. In order to validate the compatibility of GS with a well-known Golden Gate system based on standard syntax of fusion sites, we constructed a complex biosynthetic pathway using the repertoire of parts and level 1 vectors from the original MoClo system (6) and the library of GS level 2 vectors for extra flexibility. Specifically, we addressed the production of polyhydroxybutyrate (PHB) in *E. coli.* PHB is a biotechnologically useful natural polyester produced by many microorganisms under nutritional imbalances which has similar properties to conventional plastics. Production of PHB in model strains such as *Cupriavidus necator* H16 requires the action of i) 3-ketothiolase (PhaA) to condense two acetyl-CoA molecules into acetoacetyl-CoA, ii) acetoacetyl-CoA reductase (PhaB1) to reduce acetoacetyl-CoA to 3-hydroxybutyryl-CoA and iii) PHB synthase (PhaC1). To address this goal, we built the required level 0 plasmids following the procedures described by Weber (6), where: i) the Synpro16 promoter containing the AGGGGG RBS sequence (33, 34) was cloned into an A-D level 0 vector; ii) genes *phaA, phaB1*, and *phaC1* from *C. necator* H16 were domesticated, *E. coli* codon usage optimized and cloned into D-G level 0 vectors; and iii) three different terminators (*rnpB_T1, rpoC* and T500) were cloned into G-I level 0 vectors (Figure 5A). Subsequently, we assembled these level 0 parts into three level 1 plasmids using MoClo level 1 expressing *phaA*, *phaB1* and *phaC1* from positions 1, 2 and 3, respectively (see Supplementary Material, Table S3).

These level 1 plasmids were finally assembled into level 2 GS pSEVA vectors featuring different origins of replication: pBBR1, RK2 or pUC (Figure 5A). The performance of these constructs was monitored following the production of PHB in *E. coli* DH10B during 24 h by GC-MS. Considering the expected number of copies provided by each origin of replication (31), we found a positive correlation between the expected number of plasmid copies and the amount of PHB produced, i.e. 0.9931 fit (Figure 4C, Table S5). Thus, the *E. coli* strain harbouring the level 2 pUC plasmid yielded the highest PHB levels, >50% cell dry weight (CDW) followed by the strains harbouring the pBBR1 and RK2 plasmids ~13% and ~8% respectively (Figure 5B). Overall, we demonstrated that the GS system is able to incorporate level 0 parts constructed using homologous systems like CIDAR MoClo or Golden Gate while providing a compatible platform for the effective reuse and further optimization of available level 1 plasmids in the context of a broad scope of level 2 and 3 vectors with variable copy numbers.

### Golden Standard allows efficient and robust optimization of complex synthetic pathway expression using combinatorial assembly

One of the main features of MoClo assembly is the possibility to optimize the performance of a given synthetic pathway by using combinatorial assembly (35, 36). This paves the way to the construction of libraries of a given metabolic pathway under a variety of expression outputs, thus facilitating systematic exploration of expression space limited by the available parts. To accomplish this, each individual TU involved in the metabolic pathway (level 1) can be built using a set of regulatory parts, e.g. promoters, RBSs, etc. The library of individual TUs can be further combined to construct a collection of synthetic gene circuits and/or metabolic pathways (level 2). Following this approach and using a high-throughput screening method, e.g. a specific biosensor, the expression of an entire pathway can be optimized in a single step. Thus, the GS system can be used to engineer optimized gene clusters comprising several TUs. In order to validate this utility, we used GS to optimize the biosynthetic pathway for zeaxanthin production in *E. coli* (37–39). The biosynthesis of this carotenoid in *E. coli* involves five enzymatic steps: i) Synthesis of geranylgeranyl diphosphate (GGPP) from farnesyl pyrophosphate (FPP) and isopentyl diphosphate (IPP) catalysed by geranylgeranyl synthase (CrtE); ii) Condensation of two GGPP to one phytoene molecule catalysed by phytoene synthase (CrtB); iii) Desaturation of phytoene to lycopene by phytoene desaturase (CrtI); iv) Cyclization of lycopene by lycopene cyclase (CrtY); and v) Hydroxylation of β-carotene by β-carotene hydroxylase (CrtZ) (Figure 6A). The yellow colour of zeaxanthin not only simplifies the identification of colonies successfully expressing the whole pathway, but it also allows for visual recognition of relative colouration intensities between colonies. In this way, colonies expressing the optimal GS construct can be selected (11, 22, 40).

**Figure 6.**
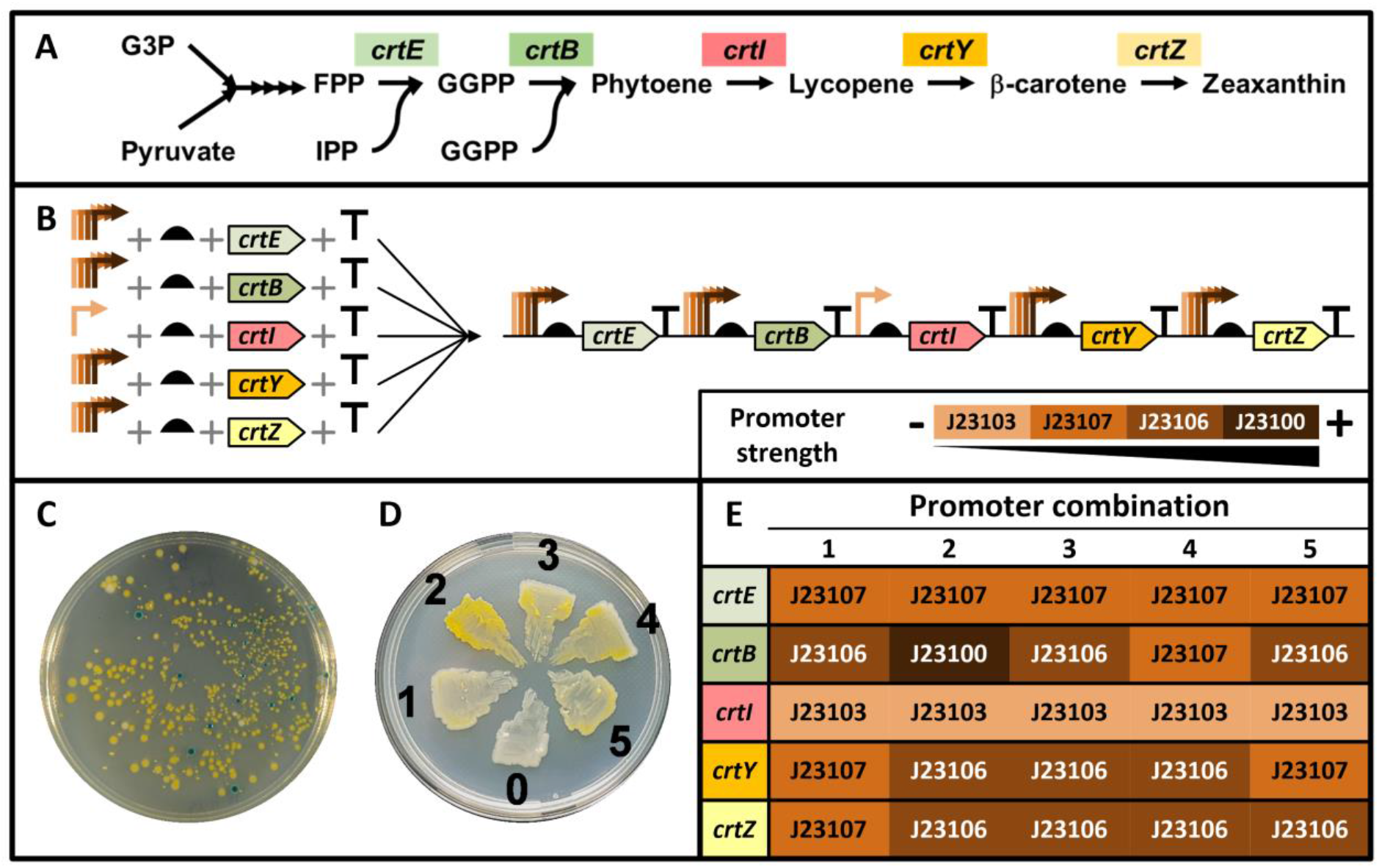
Combinatorial assembly of the zeaxanthin biosynthetic pathway using the GS system. A) Scheme of the heterologous zeaxanthin biosynthetic pathway in E. coli from glyceraldehyde 3-phosphate (G3P) and pyruvate using enzymes from P. agglomerans: FPP, farnesyl pyrophosphate; IPP, isopentyl diphosphate; GGPP, geranylgeranyl diphosphate; crtE, geranygeranyl synthase; crtB, phytoene synthase; crtI, phytoene desaturase; crtY, lycopene cyclase; crtZ, b-carotene hydroxylase. B) Scheme of the assembly of level 0 parts and level 1 TUs for zeaxanthin production with multiple level 0 Anderson promoter parts (http://parts.igem.org/Promoters/Catalog/Anderson) into a polycistronic level 2 genetic device using Golden Standard assembly. C) LB petri dish plate image showing colonies of E. coli cells transformed with the mix of level 2 parts for zeaxanthin production. Yellow colonies are positive colonies producing zeaxanthin, while the blue colonies are negative colonies carrying the empty vector. D) Restreaking of different intensity yellow colonies selected by eyesight for subsequent analysis of the promoters used (1 to 5), 0, negative control colony. E). Heat map showing the combination of Anderson’s promoters for each CDS part in the five selected colonies.

Following the procedure to construct basic level 0 parts, the five CDSs from *Pantoea agglomerans* (38) encoding the enzymes catalysing the biosynthetic pathway of zeaxanthin (*crtE, crtB, crtI, crtY* and *crtZ*), were *E. coli*-codon-usage-optimized, properly domesticated removing internal BsaI and BpiI sites and flanked by the proper D and G-based fusion sites. Subsequently, they were cloned in the SmaI site of pSEVA182. We further engineered libraries of transcription units (level 1) expressing *crtE, crtB, crtY* and *crtZ* by combinatorial assembly using an equimolar mixture of selected Anderson’s promoters (J23100, J23106, J23107 and J23116) - see http://parts.igem.org/Promoters/Catalog/Anderson, obtained from the CIDAR MoClo kit (Addgene Kit #1000000059)). To avoid the known toxicity of *crtI* overexpression (41), this gene was kept at a constant low expression level using the weak promoter J23103 (Figure 6B). Twenty positive clones (white kanamycin-resistant colonies) from each TU assembly reaction were selected, mixed and grown together. Following plasmid extractions, we generated libraries of transcription units’ level 1 plasmids for *crtE, crtB, crtY* and *crtZ* which were used to construct the synthetic level 2 cluster using a pUC-based multicopy receptor plasmid. No yellow colonies were obtained and only a few blue colonies were observed (data not shown). However, when a pBBR-based level 2 receptor plasmid was used instead, we achieved a high number of positive yellow colonies (~95%), with a transformation efficiency of 2 x 10^4^ colonies/μg of receptor plasmid (Figure 6C). Five positive colonies displaying different intensities of yellow colouration were selected for further analysis (Figure 6D). After sequencing the promoter regions, we observed a significant predominance of the medium strength promoters J23106 and J23107 (Figure 6E). It was noteworthy that colony 2, which had the most intense yellow colony colour, expressed the *crtB* gene under the control of the high strength J23100 promoter, strongly suggesting that high expression of this gene is important for a high level of zeaxanthin production. Despite the few clones analysed, it was interesting that there was an overall lack of the strong promoter J23100, which could be indicative of the toxicity of this pathway when overexpressed in *E. coli* (22, 42, 43). In fact, successful strategies for the production of carotenoids in *E. coli* generally have relied on the use of inducible promoters and/or stabilization of the pathway through chromosome integration (39, 42, 43). In this regard, the constructs for production of zeaxanthin in *E. coli* we obtained using GS combinatorial assembly, in the context of alternative copy number plasmids, highlight the usefulness of this technology in optimized synthetic metabolic pathway engineering, both in terms of improving gene expression and reducing the metabolic burden to avoid cell growth impairment and plasmid instability (44).

### Golden Standard for optimizing expression of recombinant proteins

The main applications of MoClo assembly lie in synthetic biology and focus on construction and optimization of complex metabolic pathways and/or genetic circuits featuring multiple transcription units. Applications targeting the optimization of recombinant protein production or purification are uncommon. In this context, despite MoClo having been used in modular construction of proteins including N- and/or C-terminal tags, to our knowledge this interesting feature is underexploited and lacks direct applications. Therefore, we present here a straightforward application of GS to modular construction of proteins via introduction of solubility/purification and selection/quantification tags. We used *Streptomyces antibioticus’* oleandomycin glucosyltransferase (OleD) as the model protein in our assays. This enzyme is broadly used to glycosylate many different molecules, including macrolides, flavonoids and steroids (45). In addition, crystal structures of OleD (PDB: 2IYF, 4M7P) suggest that both the N- and C-termini are likely to be located outside the globular bulk of the protein, thus making them promising candidates for modifications. We began by constructing a level 1 plasmid to support constitutive expression of OleD by the *trc* promoter and confirmed that the protein was constitutively produced in *E. coli* (Figure 7B). Furthermore, we constructed a fusion OleD protein with GFP as a C-terminal tag and confirmed that this modification did not hinder OleD enzymatic activity (Table 2). Finally, we evaluated the influence of purification/solubility tags on OleD expression and the activity of the resulting fusion proteins. To this end, we selected four commonly used N-terminal purification/solubility tags (His_6x_, strep, glutathione *S*-transferase with an HRV 3C protease cleavage site and maltose binding protein). Four GS assembly setups were prepared, one for each of the individual N-tags. Three positive clones from each GS reaction were subsequently cultivated and analysed together with clones harbouring unmodified OleD and OleD-GFP as controls (Figure 7A, see Figure S3 for details). Plasmid sequencing and SDS-PAGE analyses (Figure 7B & Figure S4) confirmed proper assembly of each of the N-terminal tags used. Results for individual MoClo assemblies are shown in Table 2 and they confirm that tags affect OleD recombinant protein production and activity in crude extracts (Figure S5).

**Figure 7.**
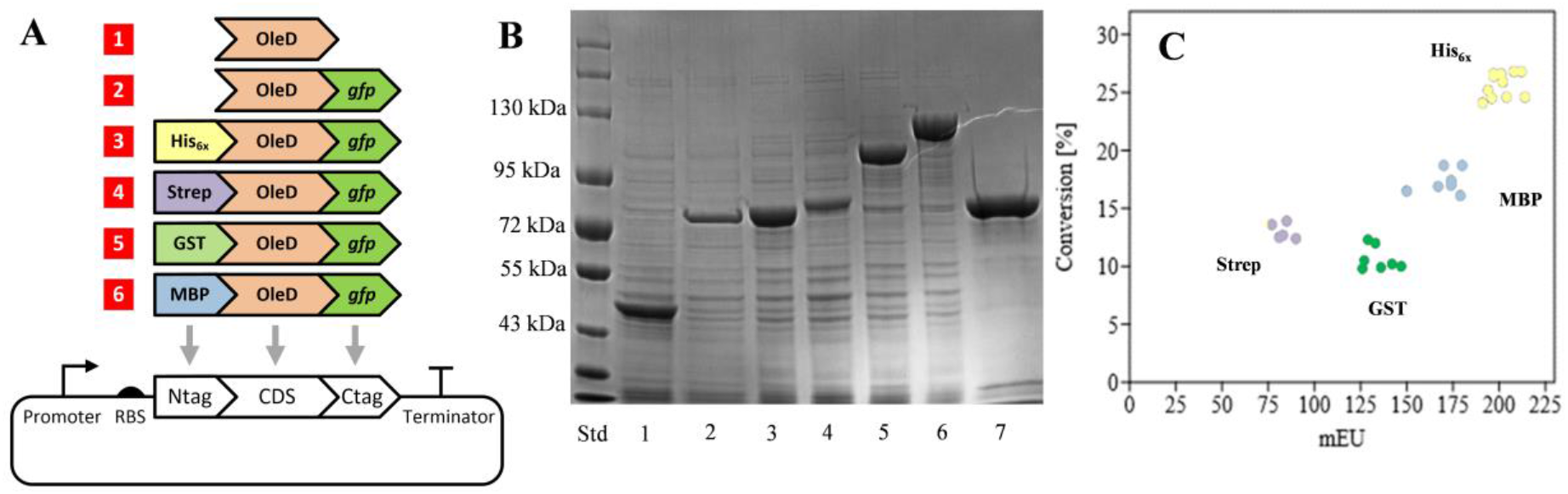
Schematic diagram and results of N- and C-terminal tag assemblies. A) Graphical representation of individual MoClo assemblies of the OleD variants tested. B) 4–20% SDS-PAGE analysis of individual assembly crude extracts. Std – NEB protein ladder (10-250 kDa); 1 – OleD; 2 – OleD-GFP; 3 – N-His-OleD-GFP; 4 – N-Strep-OleD-GFP; 5 – N-GST-OleD-GFP; 6 – N-MBP-OleD-GFP; 7 – IMAC purified N-His-OleD-GFP. C) Combined plotted results (conversion of xanthohumol to 4’-O-β-D-glucoside of xanthohumol versus fluorescence of GFP-tag) in cell-free extracts from random clones obtained using mixed MoClo assembly with different N-terminal tags.

**Table 2.** Results of individual GS assemblies of OleD with different N- or C-terminal tags. Mean values and standard deviations of three biological replicates. CFE: cell free extract. * Conversion of xanthohumol to 4’-O-β-D-glucoside of xanthohumol measured for reactions standardised on protein concentration. ** Fluorescence in cell free extracts (CFE). See Table S2 for strain descriptions.

| | Protein in CFE ($\mu\text{g mL}^{-1}$) | Conversion (%)<sup>*</sup> | Fluorescence (mEU)<sup>**</sup> |
| --- | --- | --- | --- |
| OleD (K) | 1663 ± 63 | 54 ± 1 | - |
| OleD-GFP | 1694 ± 49 | 38 ± 2 | 539 ± 16 |
| His-OleD-GFP | 1780 ± 79 | 44 ± 2 | 617 ± 27 |
| Strep-OleD-GFP | 1548 ± 101 | 20 ± 1 | 225 ± 17 |
| GST-OleD-GFP | 1859 ± 72 | 23 ± 2 | 478 ± 49 |
| MBP-OleD-GFP | 2229 ± 71 | 36 ± 2 | 553 ± 43 |

On the other hand, it is well known that protein concentration in crude extracts varies significantly and, obviously, cannot be used as the sole indicator of tag influence on activity and/or expression. Fluorescence is a more precise indicator of concentration of active GFP-tagged proteins in extracts (46), and analysis of both fluorescence and enzymatic activity is established as common practice in high-throughput screening of protein engineering libraries (47, 48). In this sense, instead of performing laborious dilutions to standardize protein concentration by means of fluorescence and further activity assays, we calculated the ratio of activity and fluorescence from the same volume of crude enzyme extracts. Taking this ratio as a key parameter, we posited that Golden Standard could be used to optimize the expression/activity of heterologously produced proteins by constructing a large combinatorial library of fusion proteins. Therefore, the four N-terminal tags were mixed stoichiometrically in the MoClo reaction and, after transformation, 30 positive clones were selected and analysed.

As expected, screening 30 random clones from the mixed assembly resulted in 4 populations that shared similar values of fluorescence and activity of cell-free extracts (CFE) (Figure 7C). Two clones from each population were sequenced to identify the tag responsible for these different behaviours. A near linear correlation between fluorescence and activity could be drawn as a logical consequence of enzyme concentration. Thus, among the different tags used, N-His_6x_ had the potential to be applied to protein purification with no negative effect on expression. On the other hand, if the aim were to screen for a protein purification tag with negligible effect on activity, the best choice would be a N-Strep-tag, which, despite displaying lowest expression levels, retained the best conversion/mEU ratio (Figure 7C). In summary, we proved that GS can be successfully applied to screening optimal recombinant protein expression. Although we present data for the constitutive expression of a recombinant protein, the same procedure could be applied to inducible promoter expression systems using a randomised library of level 1 vectors with P_T7_, P_*m*_, P_*rhaBAD*_ or any other inducible promoter. It is worth pointing out that a similar approach could be applied to serial, standardised experiments using, among others, protein linker selection, protease cleavage site selection, anchor, secretory or signal peptide selection, and could be tested in hosts other than *E. coli*.

### Golden Standard web portal

We created a website http://sysbiol.cnb.csic.es/GoldenStandard/home.php to host the Golden Standard constructions, metadata and necessary tools to design new assemblies. This is divided into multiple interconnected modules to provide flexibility for future growth. The modules are named Golden Standard DB, Golden Standard Hub and Wizard.

a. The Golden Standard DB module is the Golden Standard database interface. It features tools to search and identify DNA parts and constructions stored in the database.
b. The Golden Standard Hub module serves as a SynBioHub server (49). The data therein follows the well-established SBOL (Synthetic Biology Open Language) syntax to aid in sharing constructs. This module is linked to the Golden Standard db module.
c. The Wizard module includes a computational framework to support *in silico* assembly of constructs. This interactive tool simplifies design and assembly by allowing the user to make multiple selections of components stored in the Golden Standard Hub (levels 0, 1 or 2) and using them to simulate the cloning steps for assembly. It also supports the addition of custom elements to handle specific parts not included in the Golden Standard Hub. The user only has to provide a given sequence and its concentration to launch the simulation. The server is able to handle constructions with up to 6 level 1 elements. However, it also supports higher levels of complexity in terms of level through the use of an iterative procedure. Another key tool included in this module is the “Setup” tool, which generates a detailed report with the assembly protocol that is specific to the simulated constructs.

A step-by-step user guide to the GS Hub and Wizard is provided with the supplementary information.

## DISCUSSION AND OUTLOOK

Our use and characterization of Golden Standard assembly demonstrates that it is a robust modular cloning system capable of constructing complex expression genetic devises. Golden Standard assemblies with varying total numbers of fragments, size and DNA composition were performed to demonstrate that it is a high performance (≥95%), routine assembly method. The Golden Standard system takes advantage of: (1) a common syntax for DNA parts that is compatible with existing Golden Gate systems, thus allowing exchangeability and reusability of parts by researchers (Table 1), (2) a straightforward assembly scheme, from a single transcription unit up to genetic circuits, with a set of vectors that supports construction of up to up twenty TUs for rapid and high-throughput combinatorial assemblies, (3) a streamlined protocol supporting a logical framework and parts’ reusability for hierarchical assembly, (4) the ability to continually make improvements to the collection of building blocks (parts and modules) in order to simplify current and future engineering processes. Therefore, GS fulfils, to a great extent, the recommendations for standardization recently highlighted in the White Book on standardization in synthetic biology (50).

In order to facilitate the design of constructs using modular assembly, we developed a set of web-based tools that provide protocols and assistance for *in silico* design. We anticipate that these will enable rapid adoption of GS by non-specialist users of modular cloning. The implementation of the web server supports easy design of target synthetic pathways by: i) facilitating user-friendly selection of promoters, RBSs, and plasmid origins of replication and ii) providing detailed visualization of the final construct. Additionally, the custom-part-handling feature supports assembly procedures involving new elements and paves the way to increasing the parts database, which is expected to become a high-value asset to the scientific community.

Another advantage of the Golden Standard platform is that plasmids for genetic circuits feature a range of origins of replication to avoid undesired effects on cells. For instance, it may be necessary to use a low-to-medium copy plasmid when high expression of heterologous genetic constructs leads to an excessive metabolic burden (51) or to avoid toxic side effects on the chassis cell when membrane proteins are expressed at a high level. The comprehensive collection of parts available in the toolkit supports customized levels of gene expression.

GS is underpinned by a strong will to inspire the necessary cross-community effort to develop a universal, highly portable and standardized MoClo system. It has been designed to undergo constant updates in order to increase its functionality and scope. We envision community-guided creation of new parts to endow Golden Standard with additional modules including CRISPRi, multiplexing, genome editing, etc. Similarly, we anticipate the incorporation of quantitative outputs of parts and circuits, not only in the context of the large array of receptor vectors available, but also considering the increasing number of host environments and chassis. This feature is expected to unlock rational design and construction of complex metabolic pathways highly customized to the host chassis.

## AVAILABILITY OF MATERIALS

The plasmids listed in Table S1 have been deposited at SEVA database (http://seva-plasmids.com/) and they will be available under request. Data and tools for hierarchical assembly are available through the Golden Standard portal http://sysbiol.cnb.csic.es/GoldenStandard/home.php.

## SUPPLEMENTARY DATA

Supplementary material is provided in SI as pdf file

## AUTHORS’ CONTRIBUTIONS

BB, JTB, RK, DSL, AP and JN conceived and designed the study. BB, JTB, IM, R.K., SG, JP, SS, AW performed and analysed the experiments. DSL constructed GS web server. BB, IM, JTB DSL, JP and SS drafted the paper. EH, AP and JN: Review, editing and funding acquisition. JN Corresponding author. All authors read and approved the final manuscript.

## ACKNOWLEDGEMENTS

The authors thank Fabian Moreno, Alejandro Ronco, Darwin Carranza and Álvaro Gómez for helping in the construction of some Golden Standard plasmids. We acknowledge and appreciate plasmid donations from Till Tiso and Lars Blank (RWTH Aachen University, Aachen, Germany); Nick Wierckx (Forschungszentrum Jülich, Jülich, Germany); and Sylvestre Marillonnet (Leibniz Institute of Plant Biochemistry, Halle, Germany). The authors are specially grateful to Esteban Martínez-García and Victor de Lorenzo (Centro Nacional de Biotecnología, Consejo Superior de Investigaciones Científicas, Madrid, Spain) for their helpful, guidance and assistance adopting SEVA formalisms and critical reading of the manuscript. Finally, the authors wish to thank Clive A. Dove for proofreading of the manuscript.

