## Supplementary Materials and Methods for "Golden Standard: A complete standard, portable, and interoperative MoClo tool for model and non-model bacterial hosts"

Golden standard vectors, strains, plasmids and oligonucleotides used in this study are shown in Tables S1, S2, S3 and S4.

### Molecular biology reagents

Plasmid DNA was purified using the High Pure Plasmid Isolation Kit (Roche) and NYZminiprep (NYZTech) following the manufacturers' protocols. Genomic extractions of *C. necator* H16 were performed with the illustra bacteria genomicPrep Mini Spin Kit (GE Healthcare). DNA agarose gel bands, PCR products and digestion products were purified with illustraTM GFX PCR DNA and Gel Band Purification Kit (GE Healthcare). DNA concentration was measured using a NanoDrop 2000 Spectrophotometer (ThermoFisher Scientific). Golden Gate restriction enzymes, BpiI and BsaI-HFv2 were purchased from ThermoFisher Scientific and New England Biolabs, respectively. Phusion DNA polymerase, T4 DNA ligase and all other restriction enzymes were supplied by New England Biolabs. The MoClo toolkit was donated by Sylvestre Marillonnet via Addgene (Kit #1000000044) (1, 2).

**Construction of level 0 parts**

All level 0 parts must contain the fusion sites described in Figure 1 on the main manuscript flanked by BsaI restriction sites. Short double-stranded parts such as promoters, RBS, spacers, etc. were generated by annealing complementary pairs of oligonucleotides. Pairs of oligonucleotides were mixed to a final concentration of 1 µM with 50 mM of NaCl. Oligonucleotide mixtures were heated to 95 °C for 5 min in a thermocycler, followed by ramping down 1ºC/min per cycle for 40 cycles. These annealed parts were used for ligation into receptor vector pSEVA182 linearized by *Sma*I. The disruption of the reporter *lacZ* gene from the receptor plasmid reveals a white colony in the presence of X-gal. Positive clones were checked by sequencing and stored.

Regarding level 0 plasmid construction for PHA production, every part was PCR amplified with DNA oligonucleotides designed using Benchling (www.benchling.com) as per the following specifications: a tail containing the Bpi recognition site followed by the corresponding four nucleotide fusion site, 21 bp of minimal length for target complementarity, 50 ºC of minimal $T_m$ for that region and a maximal $T_m$ difference of ± 1.5 ºC between both oligonucleotides. The promoter was PCR-amplified from pSynPro16 (3), CDS sequences were PCR-amplified from *C. necator* H16 genomic DNA and terminator sequences were PCR-amplified from *E. coli* DH5α genome and the pBAMD1-6 plasmid (4). Primers for each PCR amplification are detailed in Table S4. Golden Gate digestion-ligation reactions were set up with 100 ng of level 0 acceptor plasmid (position AD, pICH41295; position DG, pICH41308; or position GI, pICH41276; Table S3) and the corresponding amount of gel-purified PCR product to achieve a 2:1 insert-vector molar ratio, 10 U BpiI, 400 U T4 DNA ligase and 1 mM ATP in Buffer G (Thermo Fisher Scientific; 10 mM Tris-HCl (pH 7.5 at 37°C), 10 mM $MgCl_2$, 50 mM NaCl and 0.1 mg/ml bovine serum albumin) in a reaction volume of 20 µl. The reaction was incubated in a thermocycler over four cycles of 37 °C (10 minutes) and 16 °C (10 minutes) followed by 65 °C for 20 minutes. A 100 µl aliquot of chemically competent *E. coli* DH5α was then transformed by heat shock with 5 µl of the Golden Gate reaction. Transformed cells were plated on LB-agar supplemented with 75 µg/ml streptomycin, 0.5 mM IPTG and 40 µl/ml X-gal and grown overnight at 37 °C for selection of the disruption of α-complementation β-galactosidase activity. Several white colonies per transformation were transferred to 4 mL LB medium with 75 µg/ml streptomycin and grown overnight at 37 °C for plasmid purification. The extracted plasmids were digested with BsaI to check for the presence of the correct insert size and confirmed by sequencing with primers RK81 and RK82.

#### Level 1 assemblies for PHB production

For level 1 construction of transcription units (TUs), the reaction mix contained 100 ng of acceptor plasmid (depending on the TU position, pICH47732, pICH47742, pICH47751 (Table S3) (1), a promoter plasmid part, a CDS plasmid part(s) and a terminator plasmid part in a 2:1 donor plasmid:acceptor plasmid molar ratio, 10 U BsaI, 400 U T4 DNA ligase and 1 mM ATP in Buffer G in a reaction volume of 20 µl. The reaction was incubated in a thermocycler over four: 10-minutes-at-37 °C and 10-minutes-at-16 °C cycles followed by 50 °C for 10 minutes and 80 °C for 20 minutes. Reactions were transformed into *E. coli* DH5α and plasmids prepared following the procedure described above for level 0 Golden Gate reactions, except that 100 µg/ml ampicillin, 0.5 mM IPTG and 40 µl/ml X-gal were present for selection. The extracted plasmids were digested with BbsI to check for the presence of the correct insert size and sequenced with primers RK155 and RK156 (Table S4).

#### Level 2 assemblies for PHB production

Level 2 assembly reactions were carried out with 100 ng of acceptor plasmid (pSEVA63gg19gB, pGS6219gB, and pGS6819gB; Table S1), and the plasmids with each transcription unit (pL1F1*phaC1*, pL1F2*phaA*, and pL1F3*phaB1*; Table S3) in a 2:1 donor plasmid:destination plasmid molar ratio, 10 U BpiI, 400 U T4 DNA ligase and 1 mM ATP in Buffer G in a reaction volume of 20 µl. The reaction was incubated in a thermocycler over four 10-minutes-at-37 °C and 10-minutes-at-16 °C cycles followed by 20 minutes at 65 °C. Reactions were transformed into *E. coli* DH10B following the procedure for level 0 Golden Gate reactions, except that 10 ug/ml gentamicin was added for selection. Blue-white colour selection was carried out and several white colonies were transferred to LB plates supplemented with 10 ug/mL gentamicin and 10 g/L glucose to select PHB-producing strains. Several PHB-producing colonies of each assembly were transferred to 8 mL of liquid LB medium with 10 ug/ml gentamicin for plasmid purification. The extracted plasmids were confirmed by digestion with BsaI and sequenced using primers R24, MM320, MM319, SS9, MM325, SS10, MM346 and F24 (Table S4).

**Polyhydroxybutyrate (PHB) quantification and monomer composition**

For PHB quantification (5), 25 mL of each *E. coli* strain grown in PHB production conditions for 24 hours was centrifuged for 30 minutes at 3,200 ×g. Cells were washed once with 0.85% NaCl and lyophilized for 24 hours. Lyophilized pellets were weighed to obtain the cell dry weight (CDW) of total biomass. PHB monomer composition and PHB content were determined by Gas Chromatography-Mass Spectrometry (GC-MS) of the methanolysed polyester (Table S5). 3-5 mg of lyophilized cells were suspended in 2 ml of methanol acidified with 3% (v/v) $H_2SO_4$ for PHB analysis. 2 ml of chloroform containing 0.5 mg/ml 3-methylbenzoic acid was added to the samples as an internal standard. Samples were boiled in a screw-capped tube at 100 ºC for 4 hours. After cooling, the mixture was washed twice by adding 1 ml distilled water, centrifuging for 10 minutes and removing the aqueous phase. The organic layer containing the resulting methyl ester of each monomer was analysed by GC-MS using an Agilent 7890A GC equipped with a DB-5HT capillary column (30 m length, 0.25 mm internal diameter, 0.1 µm film thickness), and mass data were collected and processed using an Agilent 5975C mass spectrometer. 1 µl samples of the organic phase were injected with helium as carrier gas at a ratio of 1:10 (1 part sample to 10 parts helium) and placed in an oven programmed to remain at 80 °C for 2 min and then increase the temperature at a rate of 5°C/min up to 115 °C for efficient peak separation. The temperature of the injector was 250 °C. Spectra were obtained as electron impacts with an ionizing energy for MS operation of 70 eV. Standard curves with known quantities of PHB (Sigma-Aldrich) dissolved in chloroform were used to calculate the monomer composition of the extracted polymers.

### Golden Standard efficiency

Efficiency of Golden Standard was estimated using a 56 bp-long spacer with fusion sites A-I to assemble level 1 constructs.

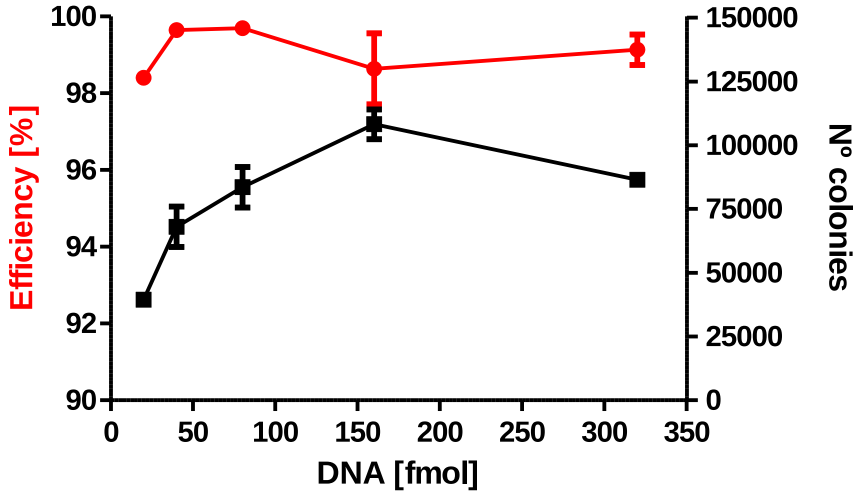


Figure S1. Golden Standard efficiency and number of colonies obtained using different DNA concentration for the assembly.

### Level 3 assembly and applications for linkers

Different linkers were constructed to develop additional applications that would otherwise require specific vectors. These linkers were short, non-coding DNA sequences flanked by BsaI or BpiI, depending on their intended application. They were named by their corresponding fusion sites following the GS rules (e.g. L_AB is a DNA linker flanked by BsaI and fusion sites A and B; L_12 is a DNA linker flanked by BpiI and fusion sites 1 and 2). Golden Standard's built-in flexibility allows users to resort to these linkers for custom applications beyond what is proposed in this paper. Furthermore, the authors encourage users to customize their own DNA constructions by including new features. Two examples related to the use of linkers are schematized in Figure S2A and 2B and explained below:

### Construction of polycistronic operons

Linkers L_AB and L_GI can be used to replace promoters and/or terminators to create polycistronic operons, respectively. These linkers are used as level 0 parts to create level 1 constructions without promoter and/or terminator. The combination of these levels 1 will generate polycistronic operons controlled by a single promoter as shown in Figure S2A.

**Construction of genetic circuits up to twenty transcription units**

To generate genetic constructions with more than six transcription units, GS provides level 3 receptor vectors that are able to integrate previously constructed level 2 circuits. The construction of a genetic circuit with a given number of transcriptional units (up to twenty) can be addressed by replacing any of the possible slots available with linkers (see Figure S2B). With this approach, there is no single rule to obtain a specific number of transcription units and both level 1 and level 2 can be replaced by their corresponding linkers. To provide a better understanding of this application, Figure S2B also shows three different ways in which a circuit comprising eleven transcription units could be built.

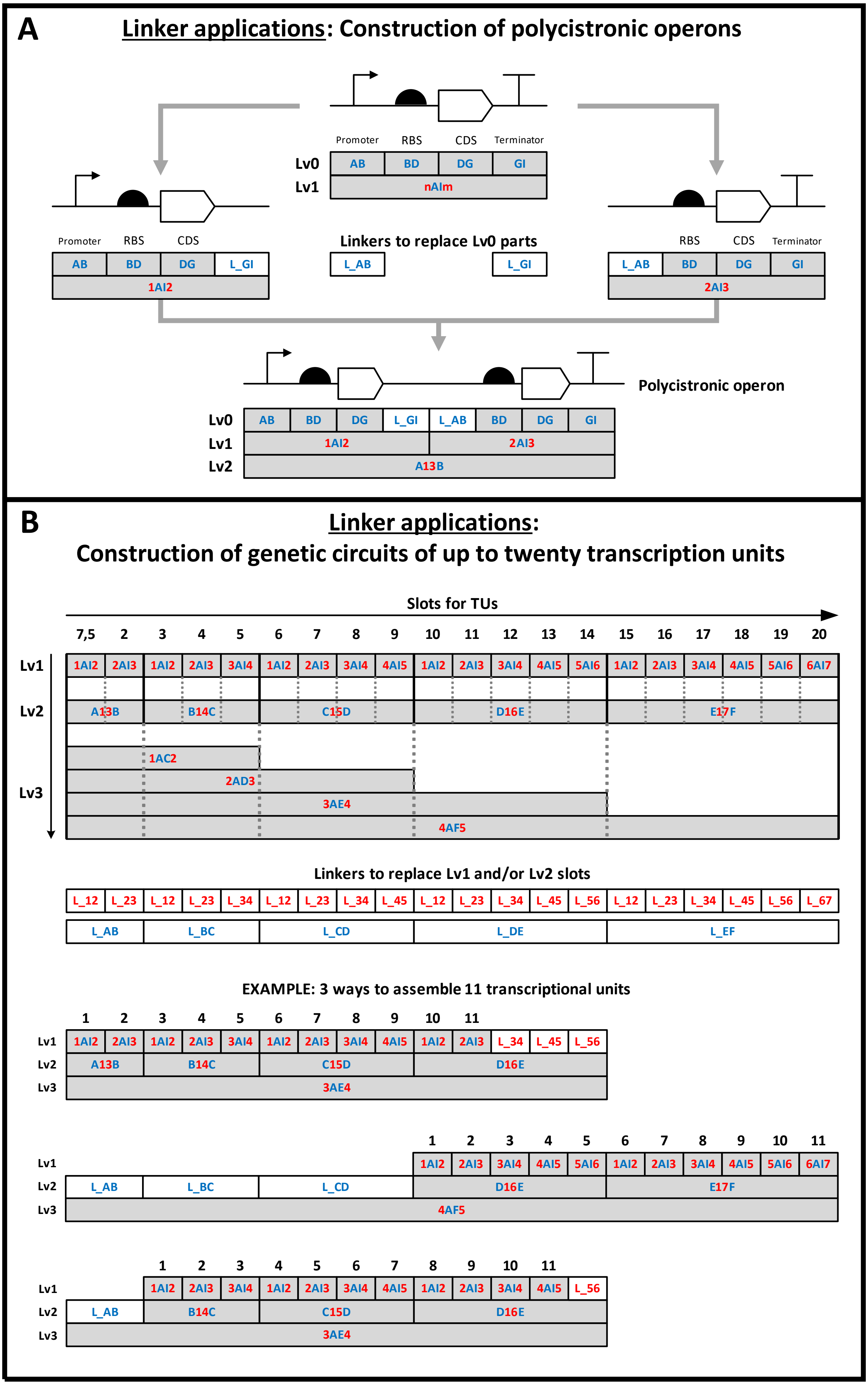
A
Linker applications: Construction of polycistronic operons
Promoter RBS CDS Terminator
Lv0 AB BD DG GI
Lv1 nAlm
Promoter RBS CDS
AB BD DG L_GI
1AI2
Linkers to replace Lv0 parts
L_AB L_GI
RBS CDS Terminator
L_AB BD DG GI
2AI3
Polycistronic operon
Lv0 AB BD DG L_GI L_AB BD DG GI
Lv1 1AI2 2AI3
Lv2 A13B
B
Linker applications:
Construction of genetic circuits of up to twenty transcription units
Slots for TUs
7,5 2 3 4 5 6 7 8 9 10 11 12 13 14 15 16 17 18 19 20
Lv1 1AI2 2AI3 1AI2 2AI3 3AI4 1AI2 2AI3 3AI4 4AI5 1AI2 2AI3 3AI4 4AI5 5AI6 1AI2 2AI3 3AI4 4AI5 5AI6 6AI7
Lv2 A13B B14C C15D D16E E17F
Lv3 1AC2 2AD3 3AE4 4AF5
Linkers to replace Lv1 and/or Lv2 slots
L_12 L_23 L_12 L_23 L_34 L_12 L_23 L_34 L_45 L_12 L_23 L_34 L_45 L_56 L_12 L_23 L_34 L_45 L_56 L_67
L_AB L_BC L_CD L_DE L_EF
EXAMPLE: 3 ways to assemble 11 transcriptional units
1 2 3 4 5 6 7 8 9 10 11
Lv1 1AI2 2AI3 1AI2 2AI3 3AI4 1AI2 2AI3 3AI4 4AI5 1AI2 2AI3 L_34 L_45 L_56
Lv2 A13B B14C C15D D16E
Lv3 3AE4
1 2 3 4 5 6 7 8 9 10 11
Lv1 1AI2 2AI3 3AI4 4AI5 5AI6 1AI2 2AI3 3AI4 4AI5 5AI6 6AI7
Lv2 L_AB L_BC L_CD D16E E17F
Lv3 4AF5
1 2 3 4 5 6 7 8 9 10 11
Lv1 1AI2 2AI3 3AI4 1AI2 2AI3 3AI4 4AI5 1AI2 2AI3 3AI4 4AI5 L_56
Lv2 L_AB B14C C15D D16E
Lv3 3AE4

Figure S2. Linker applications. A) Construction of polycistronic operons using linkers to replace promoters and/or terminators. B) Construction of genetic circuits of up to twenty transcription units using linkers to replace level 1 and/or level 2 slots.

### Use of Tags for recombinant protein expression in bacteria/Golden Standard for optimizing expression of recombinant proteins

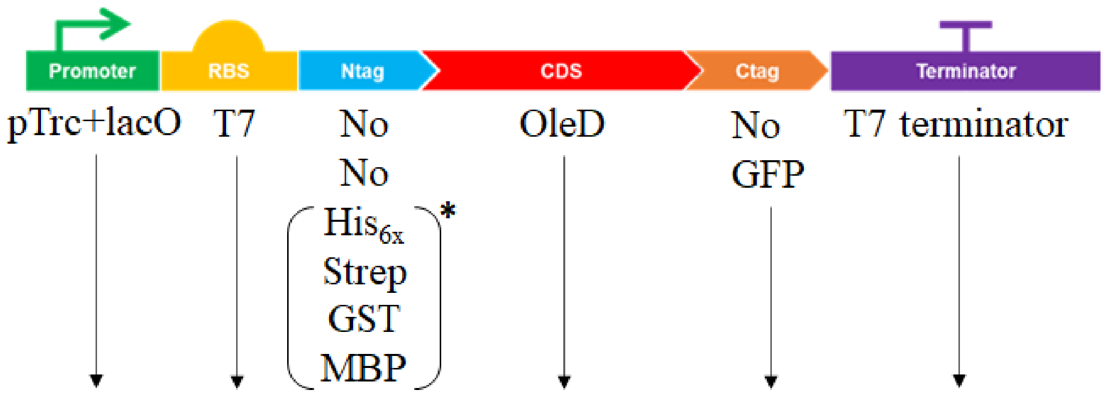


Figure S3. Visual representation of parts used in experiments with OleD glucosyltransferase. Asterisk indicates that, apart from the individual vector assemblies, all parts in brackets were used in one mixed assembly.

### SDS-PAGE

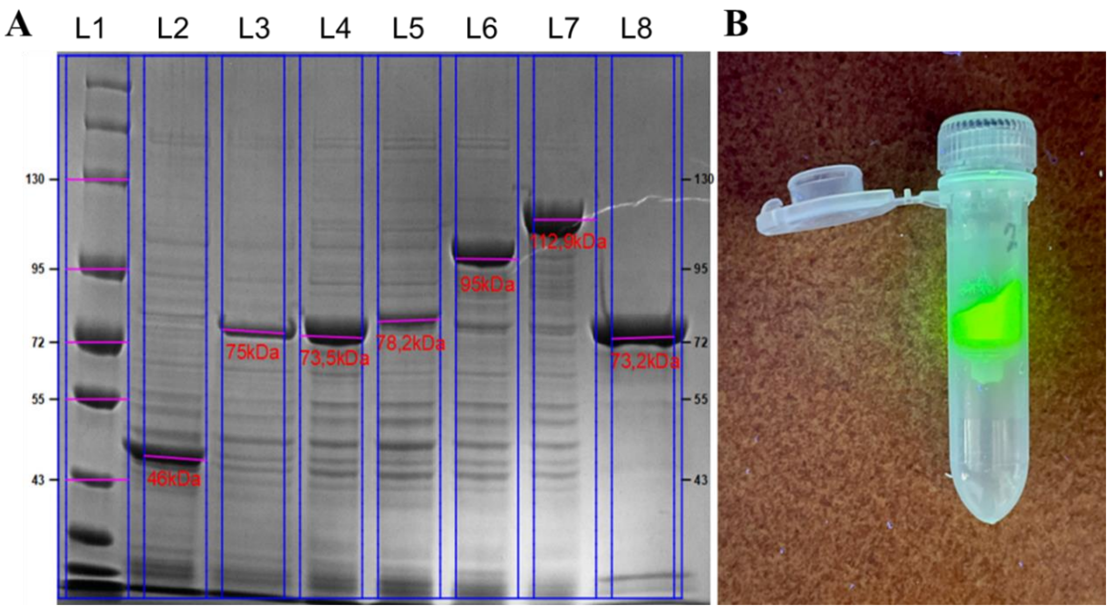

Figure S4. A) 4–20% SDS PAGE analysis of individual assembly crude extracts. L1 – NEB protein ladder (10-250 kDa); L2 – OleD; L3 – OleD-GFP; L4 – N-His-OleD-GFP; L5 – N-Strep-OleD-GFP; L6 – N-GST-OleD-GFP; L7 – N-MBP-OleD-GFP; L8 – IMAC purified N-His-OleD-GFP. B) N-His-OleD-GFP on GE Healthcare His SpinTrap column before the elution excited with UV light.

### HPLC analysis

HPLC analyses were performed using a HPLC+ Dionex Ultimate 3000 equipped with analytical C-18 Acclaim RSLC PolarAdvantage II (2.2 µm, 2.1×100 mm, Thermo Scientific) column, thermostated at 40°C set to gradient elution program with 0.1% formic acid solution in water (A) and 0.1% formic acid solution in acetonitrile (B) at 0.7 mL/min flow. From 0 to 3 min 15%->98% of eluent B, hold 98% B till 4.2 min, then from 4.2 min to 4.4 min 98%->15% of eluent B and hold till 6 min.

### Sample preparation

Glucosyltransferase reactions were stopped by addition of 100 µL of 2M HCl, followed by 350 µL of ethyl acetate. Samples were vortexed vigorously for 10 seconds, centrifuged for 1 min at 12000 x g, and then 150 µL of the organic fraction was transferred to 500 µL of methanol and analysed directly on HPLC.

### Product identification

The glucosyltransferase reaction product 4’-*O*-glucoside of xanthohumol was obtained previously and used as the reference to measure conversion (6). Measurements for conversion calculations were taken at λ=370 nm.

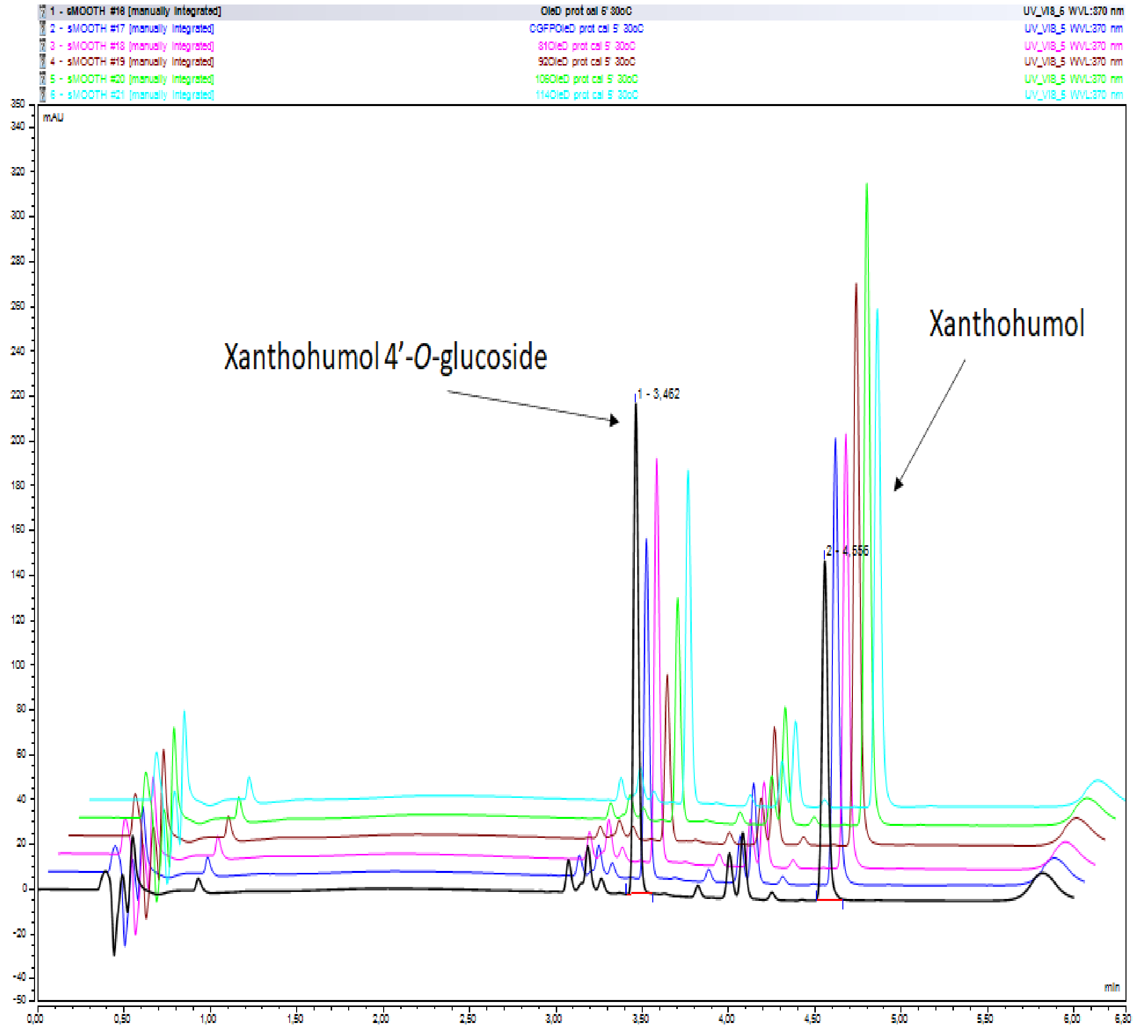


Figure S5. Example superimposed chromatograms of individual assembly reactions.

### Web portal

The website http://sysbiol.cnb.csic.es/GoldenStandard/home.php was implemented using PHP 7.0, a general-purpose scripting language suited to web development. The relational database management system MariaDB server 10.4.18 was chosen to store the schema and data for the constructions and experiment metadata. The rest of software tools included in the portal were developed using Javascript and PHP. The Golden Standard DB module is a forked version of SynBioHub, a web-based repository for synthetic biology typically used to share synthetic biology designs. The SubmitSnapgene and DownloadSnapgene plugins were included to provide extra flexibility to upload other types of construction files, whereas Visual ComponentUse and VisualSeqviz improve graph quality.

### Golden Standard wizard guide

To access the Golden Standard wizard, users must click on the ***Golden Standard wizard*** button featured in the landing page.

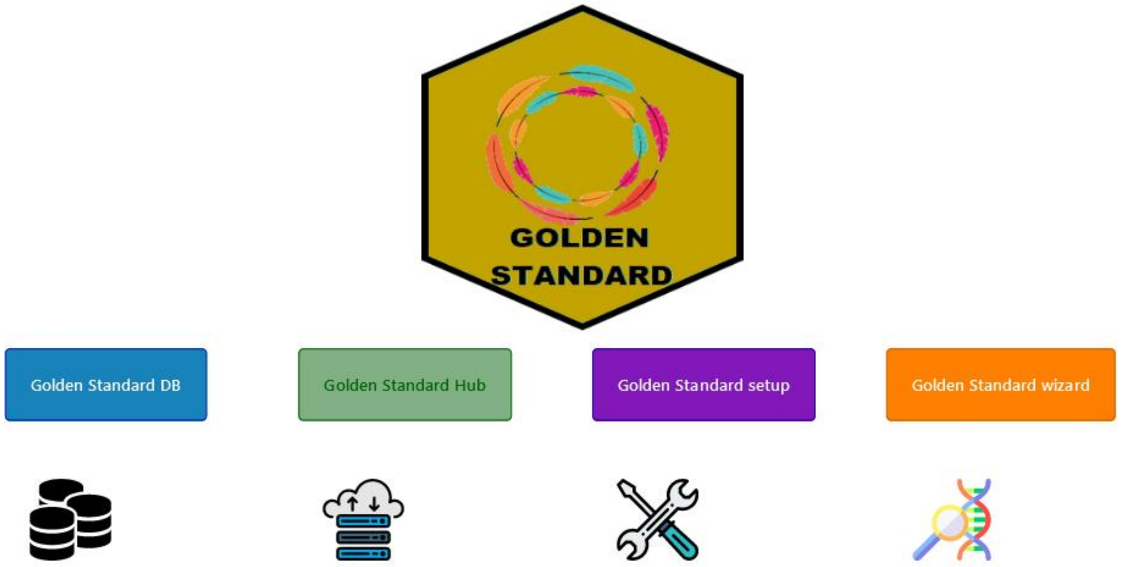


This opens a new page with the multiple options. To start a new assembly:

1. Click on 1AI2 construct green button.

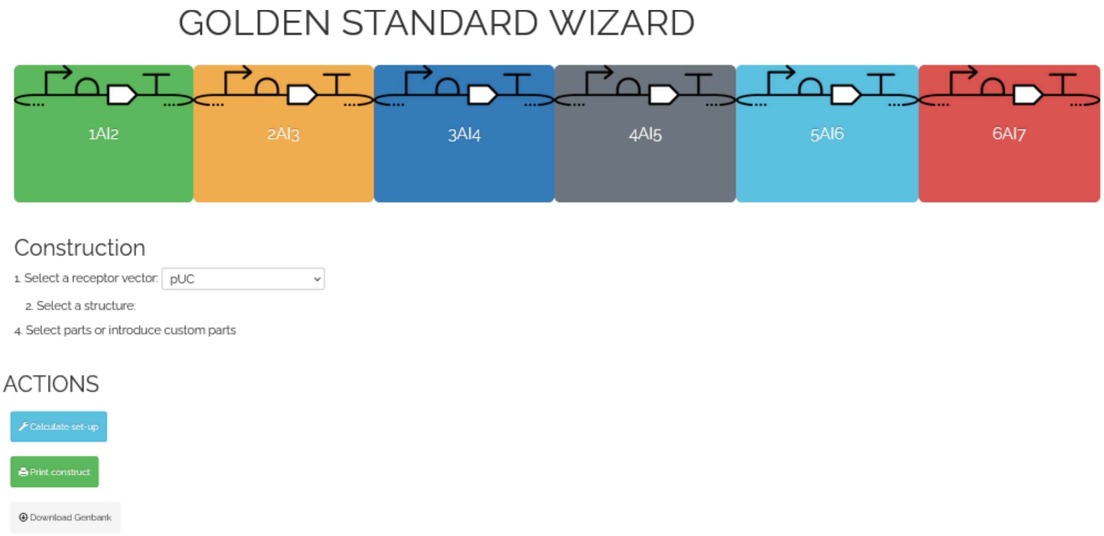


2. A form is deployed with the basic structure of level 1 constructs and options to enter information:

    1. The first step is to select a receptor vector.

2. The structure can be selected with or without C- terminal or N-terminal tags based on Golden Standard toolkit restrictions. To select an option, click on the radio buttons. The predicted structure will appear in the bottom figure.

3. Orientation can be modified selecting **Forward** or **Reverse.**

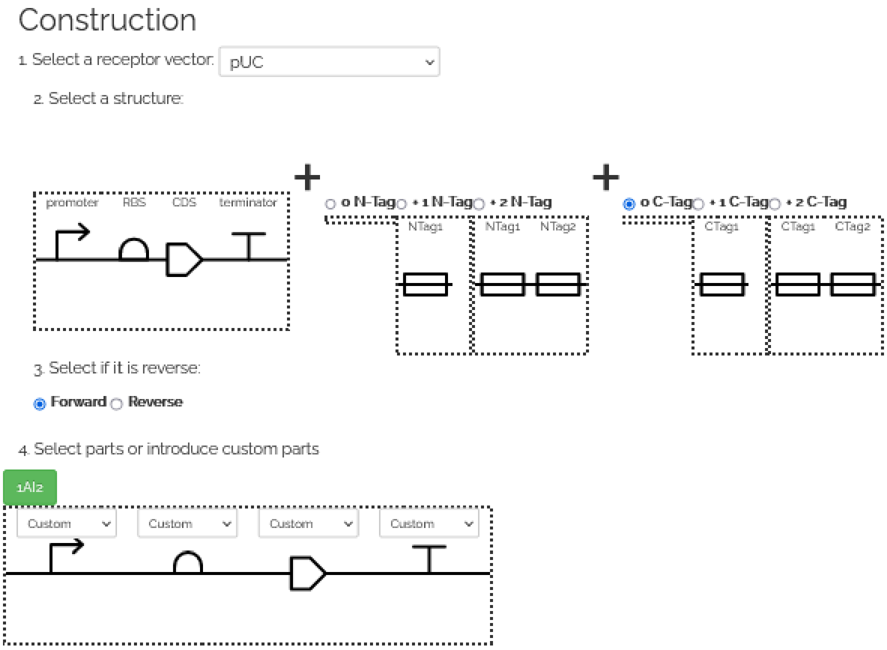


4. Next it is possible to select the parts. The bottom figure (above the parts symbols) features combo boxes with the available parts. For instance, clicking the promoter combo box will open up the list of available promoters.

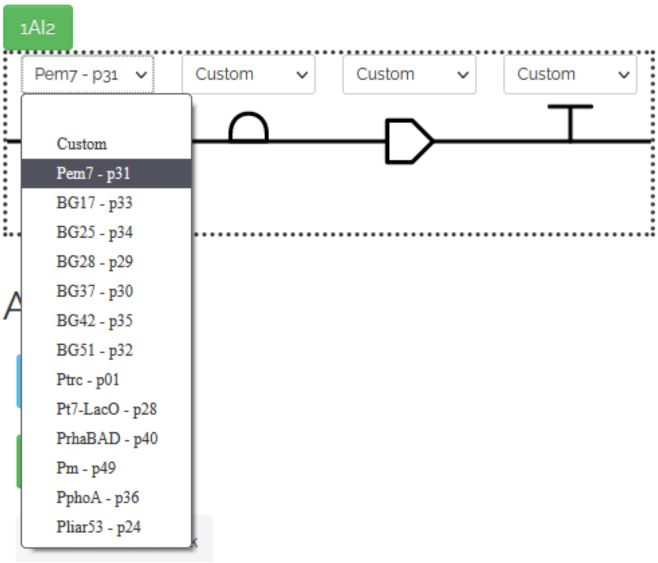

5. Select a promoter.

   A. To use a promoter already integrated in Golden Standard, select an element with an identification tag from the list.

      1. The id will appear on the promoter icon.

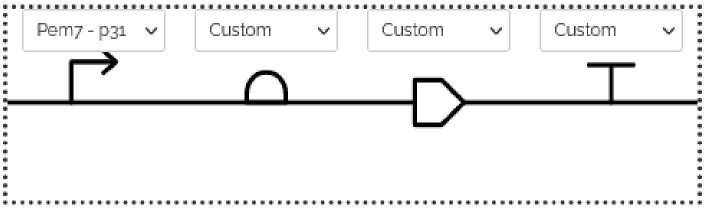


   B. To use a custom promoter, follow these steps:

      1. Click on the combo box and select Custom from the drop-down list.

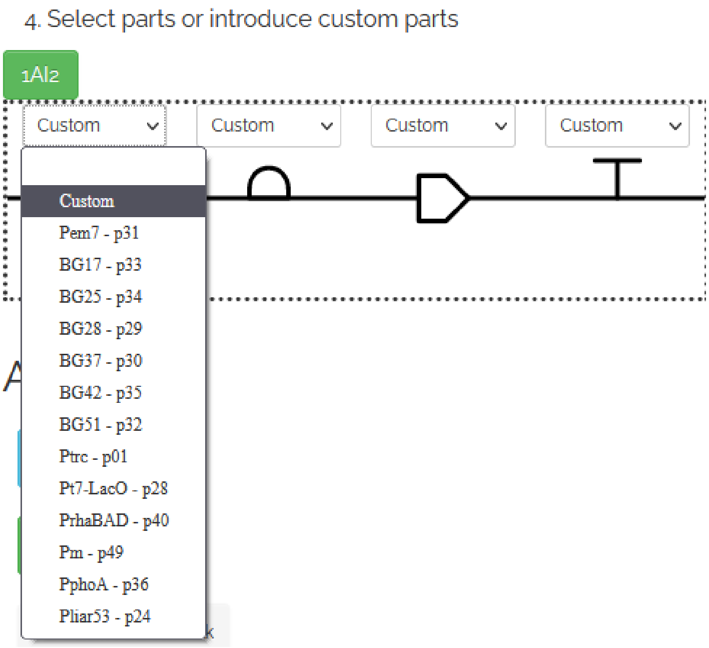


      2. A new panel opens up prompting the user to enter the name, sequence and concentration of the new promoter.

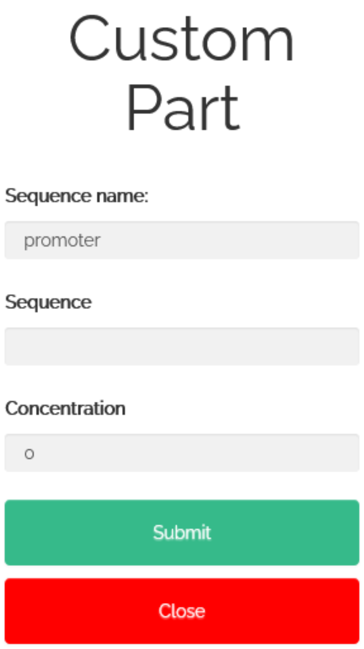


3. Enter the required information: ID, sequence and concentration (optional) and click on Submit.

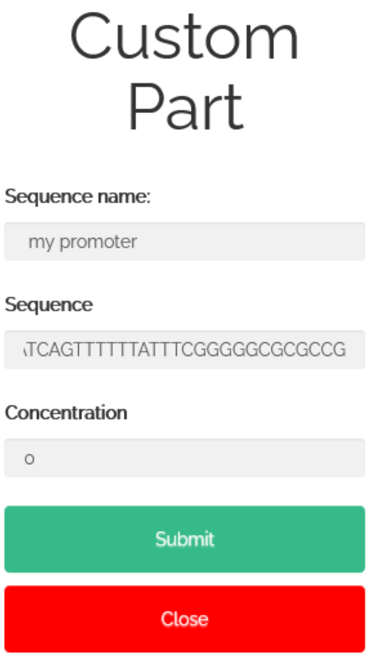

6. Select an RBS following the same procedure. To create a custom element follow steps 5.B 1-3.

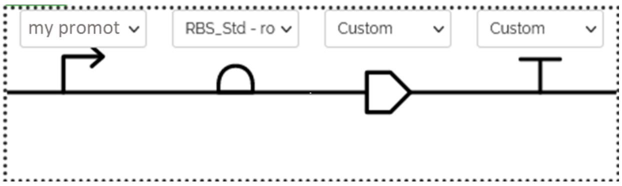


7. Select a CDS from the library or create a CDS. To create a custom element follow steps 5.B 1-3.

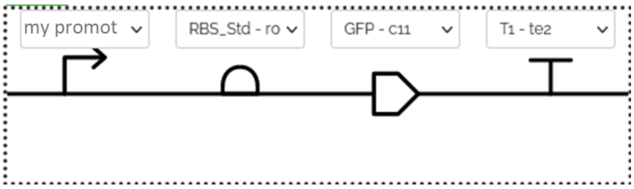


8. Finally, the last step is the terminator selection. To create a custom element follow steps 5.B 1-3.

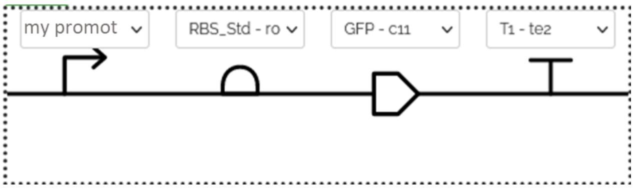


9. Next it is possible to select a new construct, for instance 2AI3 and repeat steps 3 to 9.

10. The process is identical for 3AI4, 4AI5, 5AI6 and 6AI7.

11. To visualize the result, click on Download Genbank. This will download a Genbank sequence format with the *in silico* assembly.

12. To complete the protocol with TU concentrations, click on Calculate set-up and a new window will open.

| Tube label | TU | Id | Conc.(ng/ul) | Size(bp) | | ul to get final volume (fmol) |
|---|---|---|---|---|---|---|
| Tube label | 1AI2 | Id | 0 | 0 | | 0 |
| | 2AI3 | Id | 0 | 0 | | 0 |
| | 3AI4 | Id | 0 | 0 | | 0 |
| | 4AI5 | Id | 0 | 0 | | 0 |
| | 5AI6 | Id | 0 | 0 | | 0 |
| | 6AI7 | Id | 0 | 0 | | 0 |
| | receptor_vector | Id | 0 | 0 | | 0 |
| | **Enzyme** | | | | | 0,5 |
| | **Ligase** | | | | | 0,5 |
| | **Water** | | | | | 17 |
| | **Buffer** | | | | | 2 |
| | | | | **Subtotal** | 0 | |
| | | | | | **Total** | 0 |

13. By clicking on each of the boxes it is possible to enter the required values to set up an experiment and calculate the amount of water required.

Table S1. List of Golden Standard system vectors available.

| Well | Plasmid name | Level | Antib. | OriV | Purpose |
|---|---|---|---|---|---|
| **A1** | pSEVA182 | - | Amp | pUC | Lv 0 part receptor |
| **A2** | pSEVA23g19g1 | 1 | Km | pBBR | Host vector Lv1, 1AI2 cargo |
| **A3** | pSEVA23g19g2 | 1 | Km | pBBR | Host vector Lv1, 2AI3 cargo |
| **A4** | pSEVA23g19g3 | 1 | Km | pBBR | Host vector Lv1, 3AI4 cargo |
| **A5** | pSEVA23g19g4 | 1 | Km | pBBR | Host vector Lv1, 4AI5 cargo |
| **A6** | pSEVA23g19g5 | 1 | Km | pBBR | Host vector Lv1, 5AI6 cargo |
| **A7** | pSEVA23g19g6 | 1 | Km | pBBR | Host vector Lv1, 6AI7 cargo |
| **A8** | pSEVA23g19g1R | 1 | Km | pBBR | Host vector Lv1, 1IA2 cargo |
| **A9** | pSEVA2819g1 | 1 | Km | pUC | Host vector Lv1, 1AI2 cargo |
| **A10** | pSEVA2819g2 | 1 | Km | pUC | Host vector Lv1, 2AI3 cargo |
| **A11** | pSEVA2819g3 | 1 | Km | pUC | Host vector Lv1, 3AI4 cargo |
| **A12** | pSEVA2819g4 | 1 | Km | pUC | Host vector Lv1, 4AI5 cargo |
| **B1** | pSEVA2819g5 | 1 | Km | pUC | Host vector Lv1, 5AI6 cargo |
| **B2** | pSEVA2819g6 | 1 | Km | pUC | Host vector Lv1, 6AI7 cargo |
| **B3** | pSEVA2819g1R | 1 | Km | pUC | Host vector Lv1, 1IA2 cargo |
| **B4** | pSEVA2219g1 | 1 | Km | RK2 | Host vector Lv1, 1AI2 cargo |
| **B5** | pSEVA2219g2 | 1 | Km | RK2 | Host vector Lv1, 2AI3 cargo |
| **B6** | pSEVA2219g3 | 1 | Km | RK2 | Host vector Lv1, 3AI4 cargo |
| **B7** | pSEVA2219g4 | 1 | Km | RK2 | Host vector Lv1, 4AI5 cargo |
| **B8** | pSEVA2219g5 | 1 | Km | RK2 | Host vector Lv1, 5AI6 cargo |
| **B9** | pSEVA2219g6 | 1 | Km | RK2 | Host vector Lv1, 6AI7 cargo |
| **B10** | pSEVA2219g1R | 1 | Km | RK2 | Host vector Lv1, 1IA2 cargo |
| **B11** | pSEVA63g19gA | 2 | Gm | pBBR | Host vector Lv2, 1CD4 cargo. 2 TUs |
| **B12** | pSEVA63g19gB | 2 | Gm | pBBR | Host vector Lv2, B14C cargo. 3 TUs |
| **C1** | pSEVA63g19gC | 2 | Gm | pBBR | Host vector Lv2, C15D cargo. 4 TUs |
| **C2** | pSEVA63g19gD | 2 | Gm | pBBR | Host vector Lv2, D16E cargo. 5 TUs |
| **C3** | pSEVA63g19gE | 2 | Gm | pBBR | Host vector Lv2, E17F cargo. 6 TUs |
| **C4** | pSEVA6819gA | 2 | Gm | pUC | Host vector Lv2, 1CD4 cargo. 2 TUs |
| **C5** | pSEVA6819gB | 2 | Gm | pUC | Host vector Lv2, B14C cargo. 3 TUs |
| **C6** | pSEVA6819gC | 2 | Gm | pUC | Host vector Lv2, C15D cargo. 4 TUs |
| **C7** | pSEVA6819gD | 2 | Gm | pUC | Host vector Lv2, D16E cargo. 5 TUs |
| **C8** | pSEVA6819gE | 2 | Gm | pUC | Host vector Lv2, E17F cargo. 6 TUs |
| **C9** | pSEVA6219gA | 2 | Gm | RK2 | Host vector Lv2, 1CD4 cargo. 2 TUs |
| **C10** | pSEVA6219gB | 2 | Gm | RK2 | Host vector Lv2, B14C cargo. 3 TUs |
| **C11** | pSEVA6219gC | 2 | Gm | RK2 | Host vector Lv2, C15D cargo. 4 TUs |
| **C12** | pSEVA6219gD | 2 | Gm | RK2 | Host vector Lv2, D16E cargo. 5 TUs |
| **D1** | pSEVA6219gE | 2 | Gm | RK2 | Host vector Lv2, E17F cargo. 6 TUs |
| **D2** | pSEVA23g19gα | 3 | Km | pBBR | Host vector Lv3, 1AC2 cargo. 5 TUs |
| **D3** | pSEVA23g19gβ | 3 | Km | pBBR | Host vector Lv3, 2AD3 cargo. 9 TUs |
| **D4** | pSEVA23g19gγ | 3 | Km | pBBR | Host vector Lv3, 3AE4 cargo. 14 TUs |
| **D5** | pSEVA23g19gδ | 3 | Km | pBBR | Host vector Lv3, 4AF5 cargo. 20 TUs |
| **D6** | pSEVA182-Pem7_AB | 0 | Amp | pUC | Pem7 constitutive promoter |
| **D7** | pSEVA182-BG17_AB | 0 | Amp | pUC | BG17 constitutive promoter adapted from Zobel et al. (7) |
| **D8** | pSEVA182-BG25_AB | 0 | Amp | pUC | BG25 constitutive promoter adapted from Zobel et al. (7) |

| | | | | | |
|---|---|---|---|---|---|
| **D9** | pSEVA182-BG28_AB | 0 | Amp | pUC | BG28 constitutive promoter adapted from Zobel et al. (7) |
| **D10** | pSEVA182-BG37_AB | 0 | Amp | pUC | BG37 constitutive promoter adapted from Zobel et al. (7) |
| **D11** | pSEVA182-BG42_AB | 0 | Amp | pUC | BG42 constitutive promoter adapted from Zobel et al. (7) |
| **D12** | pSEVA182-BG51_AB | 0 | Amp | pUC | BG51 constitutive promoter adapted from Zobel et al. (7) |
| **E1** | pSEVA18-Ptrc_AB | 0 | Amp | pUC | Ptrc promoter repressed by LacI |
| **E2** | pSEVA182-Pt7-LacO_AB | 0 | Amp | pUC | Pt7-LacO promoter repressed by LacI |
| **E3** | pSEVA182-PrhaBAD_AB | 0 | Amp | pUC | PrhaBAD promoter activated by RhaRS protein and L-rhamnose |
| **E4** | pSEVA182-Pm_AB | 0 | Amp | pUC | Pm promoter activated by XylS protein and 3-methylbenzoate |
| **E5** | pSEVA182-PphoA_AB | 0 | Amp | pUC | PphoA promoter repressed by phosphate from Torres-Bacete *et al*. (8) |
| **E6** | pSEVA182-Pliar53_AB | 0 | Amp | pUC | Pliar promoter repressed by phosphate from Torres-Bacete *et al*. (8) |
| **E7** | pSEVA182-Std_BD | 0 | Amp | pUC | RBS consensus adapted from Nogales *et al*. (9) |
| **E8** | pSEVA182-T7RBS_BD | 0 | Amp | pUC | RBS from pET plasmid |
| **E9** | BCD12_AB | 0 | Amp | pUC | Bicistronic RBS. Medium strenght |
| **E10** | BCD2_AB | 0 | Amp | pUC | Bicistronic RBS. High strenght |
| **E11** | B0034_AB | 0 | Amp | pUC | Monocistronic RBS. High strenght |
| **E12** | pSEVA-T7RBS_BC | 0 | Amp | pUC | RBS from pET plasmid |
| **F1** | pSEVA182-GFP_DG | 0 | Amp | pUC | Green Fluorescent gene |
| **F2** | pSEVA182-RFP_DG | 0 | Amp | pUC | Red Fluorescent gene |
| **F3** | pSEVA182-YFP_DG | 0 | Amp | pUC | Yellow Fluorescent gene |
| **F4** | pSEVA182-BFP_DG | 0 | Amp | pUC | Blue Fluorescent gene |
| **F5** | pSEVA182-T1_GI | 0 | Sm/Sp | pUC | T1 terminator from *rnpB* gene of E. coli MG1655 |
| **F6** | pSEVA182-rpoC_GI | 0 | Sm/Sp | pUC | Terminator from *rpoBC* operon of E. coli |
| **F7** | pSEVA182-T500_GI | 0 | Sm/Sp | pUC | T500 Terminator |
| **F8** | pSEVA182-Bba_B0014_GI | 0 | Amp | pUC | Bba_B1006 Artificial terminator |
| **F9** | pSEVA182-Bba_B0014_HI | 0 | Amp | | Bba_B1006 Artificial terminator |
| **F10** | pSEVA182-N-6xHis-tag_CD | 0 | Amp | pUC | N-6xHis tag |
| **F11** | pSEVA182-N-Strep-tag_CD | 0 | Amp | pUC | N-Strep tag |
| **F12** | N-GST HRV3C tag_CD | 0 | Amp | Col E1 | N-GST HRV3C tag |

| | | | | | |
|---|---|---|---|---|---|
| **G1** | N-MBP tag_CD | 0 | Amp | Col E1 | N-MBP tag |
| **G2** | pSEVA182-N-OmpA-tag_CD | 0 | Amp | pUC | N-OmpA tag |
| **G3** | pSEVA182-N-PelB_CD | 0 | Amp | pUC | N-PelB |
| **G4** | pSEVA182-C-6xHis-tag_FG | 0 | Amp | pUC | C-6xHistag |
| **G5** | pSEVA182-C-Streptag_FG | 0 | Amp | pUC | C-Streptag |
| **G6** | pSEVA182-C-LinkGFP_FG | 0 | Amp | pUC | C-LinkGFP |
| **G7** | pSEVA181-L_12 | 0 | Amp | pUC | Linker with 12 fusion site (BsaI) |
| **G8** | pSEVA181-L_23 | 0 | Amp | pUC | Linker with 23 fusion site (BsaI) |
| **G9** | pSEVA181-L_34 | 0 | Amp | pUC | Linker with 34 fusion site (BsaI) |
| **G10** | pSEVA181-L_45 | 0 | Amp | pUC | Linker with 45 fusion site (BsaI) |
| **G11** | pSEVA181-L_56 | 0 | Amp | pUC | Linker with 56 fusion site (BsaI) |
| **G12** | pSEVA181-L_67 | 0 | Amp | pUC | Linker with 67 fusion site (BsaI) |
| **H1** | pSEVA182-L_AB | 0 | Amp | pUC | Linker with AB fusion site (BpiI) |
| **H2** | pSEVA182-L_BC | 0 | Amp | pUC | Linker with BC fusion site (BpiI) |
| **H3** | pSEVA182-L_CD | 0 | Amp | pUC | Linker with CD fusion site (BpiI) |
| **H4** | pSEVA182-L-DE | 0 | Amp | pUC | Linker with DE fusion site (BpiI) |
| **H5** | pSEVA182-L-EF | 0 | Amp | pUC | Linker with EF fusion site (BpiI) |
| **H6** | pSEVA182-L-FG | 0 | Amp | pUC | Linker with FG fusion site (BpiI) |
| **H7** | pSEVA182-L-GH | 0 | Amp | pUC | Linker with GH fusion site (BpiI) |
| **H8** | pSEVA182-L-HI | 0 | Amp | pUC | Linker with HI fusion site (BpiI) |
| **H9** | pSEVA182-L-GI | 0 | Amp | pUC | Linker with GI fusion site (BpiI) |
| **H10** | pSEVA28-xylS_12 | 1 | Km | pUC | XylS level 1 at position 12 |
| **H1** | pSEVA23g-rhaRS_12 | 1 | Km | pBBR | RhaRS level 1 at position 12 |
| **H12** | pSEVA23g-lacI_12 | 1 | Km | pBBR | LacI level 1 at position 12 |

Table S2. List of strains used in this study.

| Strain | Genotype | Ref |
|---|---|---|
| ***E. coli* DH5α** | fhuA2 Δ(argF-lacZ)U169 phoA glnV44 Φ80 Δ(lacZ)M15 gyrA96 recA1 relA1 endA1 thi-1 hsdR17 | New England Biolabs |
| ***E. coli* DH10B** | Δ*(ara-leu)7697 araD139 tonA* Δ*lacX74* duplication(514341-627601) *galK16 galE15 e14- ϕ80dlacZ*Δ*M15 recA1 relA1 endA1* Tn*10.10 nupG rpsL*(Str$^R$) *rph spoT1* Δ*(mrr-hsdRMS-mcrBC)* | (10) |
| ***P. putida* KT2440** | Wild-type strain derived of P. putida mt-2 cured of the pWW0 plasmid | (11) |
| ***E. coli* DH5α pSEVA2219g2-pem7stGFP** | pSEVA2219g2-pem7stGFP plasmid | This work |
| ***E. coli* DH5α pSEVA23g19g2-pem7stGFP** | pSEVA23g19g2-pem7stGFP plasmid | This work |
| ***E. coli* DH5α** | pSEVA2819g2-pem7stGFP plasmid | This work |

| | | |
|---|---|---|
| **pSEVA2819g2-pem7stGFP** | | |
| ***P. putida* KT2440 pSEVA2219g2-pem7stGFP** | pSEVA2219g2-pem7stGFP plasmid | This work |
| ***P. putida* KT2440 pSEVA23g19g2-pem7stGFP** | pSEVA23g19g2-pem7stGFP plasmid | This work |
| ***E. coli* DH5α Lv1 crtE** | Collection of level 1 crtE vectors | This work |
| ***E. coli* DH5α Lv1 crtB** | Collection of level 1 crtB vectors | This work |
| ***E. coli* DH5α Lv1 crtI** | Collection of level 1 crtI vectors | This work |
| ***E. coli* DH5α Lv1 crtY** | Collection of level 1 crtY vectors | This work |
| ***E. coli* DH5α Lv1 crtZ** | Collection of level 1 crtZ vectors | This work |
| ***E. coli* DH5α Lv2 zeaxanthin pUC** | Level 2 zeaxanthin plasmid with pUC origin of replication | This work |
| ***E. coli* DH5α Lv2 zeaxanthin pBBR** | Level 2 zeaxanthin plasmid with pBBR1 origin of replication | This work |
| ***E. coli* DH5α pL1F1-*phaC1*** | pL1F1-*phaC1* plasmid | This work |
| ***E. coli* DH5α pL1F2-*phaA*** | pL1F2-*phaA* plasmid | This work |
| ***E. coli* DH5α pL1F3-*phaB1*** | pL1F3-*phaB1* plasmid | This work |
| ***E. coli* DH10B pSEVA63g19gB-PHB** | pSEVA63g19gB-PHB plasmid with pBBR1 origin of replication | This work |
| ***E. coli* DH10B pSEVA6219gB-PHB** | pSEVA6219gB-PHB plasmid with RK2 origin of replication | This work |
| ***E. coli* DH10B pSEVA6819gB-PHB** | pSEVA6819gB-PHB plasmid with pUC origin of replication | This work |
| ***E. coli* DH5α Lv 1 OleD** | Level 1 OleD plasmid | This work |
| ***E. coli* DH5α Lv1 OleD-C-GFP** | Level 1 OleD-C-GFP plasmid | This work |
| ***E. coli* DH5α Lv1 N-His-OleD-C-GFP** | Level 1 N-His-OleD-C-GFP plasmid | This work |
| ***E. coli* DH5α Lv1 N-Strep-OleD-C-GFP** | Level 1 N-Strep-OleD-C-GFP plasmid | This work |
| ***E. coli* DH5α Lv1 N-GST-OleD-C-GFP** | Level 1 N-GST-OleD-C-GFP plasmid | This work |
| ***E. coli* DH5α Lv1 N-MBP-OleD-C-GFP** | Level 1 N-MBP-OleD-C-GFP plasmid | This work |

Table S3. List of plasmids used in this study.

| Plasmid | Characteristics | Ref |
|---|---|---|
| **pSEVA182-Pem7_AB** | $Amp^R$, pUC, Lv0 promoter Pem7, AB fusion sites | This work |
| **J23100_AB** | $Amp^R$, pUC, Lv0 promoter J23100, AB fusion sites | (12) |
| **J23106_AB** | $Amp^R$, pUC, Lv0 promoter J23106, AB fusion sites | (12) |
| **J23107_AB** | $Amp^R$, pUC, Lv0 promoter J23107, AB fusion sites | (12) |
| **J23116_AB** | $Amp^R$, pUC, Lv0 promoter J23116, AB fusion sites | (12) |

| | | |
|---|---|---|
| **J23103_AB** | Amp$^R$, pUC, Lv0 promoter J23103, AB fusion sites | (12) |
| **pSEVA18-PTrc-LacO_AB** | Amp$^R$, pUC, Lv0 Trc strong promoter with LacO, AB fusion sites | This work |
| **pRK154** | Sm$^R$, pMB1/ColE2, Lv0 λT0 terminator with SynPro16 promoter, AD fusion sites | This work |
| **pSEVA182-RBS-ST_BD** | Amp$^R$, pUC, Lv0 consensus RBS standard, BD fusion sites | This work |
| **pSEVA18-RBS-T7_BD** | Amp$^R$, pUC, Lv0 strong RBS from pET28, BD fusion sites | This work |
| **pSEVA18-RBS-T7_BC** | Amp$^R$, pUC, Lv0 strong RBS from pET28, BC fusion sites | This work |
| **pSEVA18-6xHis Tag-_CD** | Amp$^R$, pUC, Lv0 6xHis tag, CD fusion sites | This work |
| **pSEVA18-Strep Tag-_CD** | Amp$^R$, pUC, Lv0 Strep, CD fusion sites | This work |
| **pSEVA18-GST Tag-_CD** | Amp$^R$, pUC, Lv0 GST with HRV 3C protease clavage site, CD fusion sites | This work |
| **pSEVA18-MBP Tag-_CD** | Amp$^R$, pUC, Lv0 MBP tag, CD fusion sites | This work |
| **pSEVA182-GFP_DG** | Amp$^R$, pUC, Lv0 *gfp* gene, DG fusion sites | This work |
| **p181-crtE_DG** | Amp$^R$, pUC, Lv0 *crtE* gene, DG fusion sites | This work |
| **p181-crtB_DG** | Amp$^R$, pUC, Lv0 *crtB* gene, DG fusion sites | This work |
| **p181-crtI_DG** | Amp$^R$, pUC, Lv0 *crtI* gene, DG fusion sites | This work |
| **p181-crtY_DG** | Amp$^R$, pUC, Lv0 *crtY* gene, DG fusion sites | This work |
| **p181-crtZ_DG** | Amp$^R$, pUC, Lv0 *crtZ* gene, DG fusion sites | This work |
| **pRK208-*phaA*_DG** | Sm$^R$, pUC, Lv0 *phaA* gene, DG fusion sites | This work |
| **pRK209-*phaB1*_DG** | Sm$^R$, pUC, Lv0 *phaB1* gene, DG fusion sites | This work |
| **pRK210-*phaC*_DG** | Sm$^R$, pUC, Lv0 *phaC* gene, DG fusion sites | This work |
| **pMA-T-OleD_DG** | Amp$^R$, pUC, Lv0 *OleD* gene, DG fusion sites | This work |
| **pSEVA18-GFP Tag-_FG** | Amp$^R$, pUC, Lv0 GFP gene with N-terminal TSAAA linker, FG fusion sites | This work |
| **pSEVA18-T7-Terminator_GI** | Amp$^R$, pUC, Lv0 strong Terminator from pET28, GI fusion sites | This work |
| **pSEVA182-BBa_B1006_GI** | Amp$^R$, ori pUC, Lv0 BBa_B1006 terminator, GI fusion sites | This work |
| **pRK107-rpoC_GI** | Sm$^R$, pUC, Lv0 GGtλT0-L0 terminator, GI fusion sites | This work |
| **pRK34-GGtλT0-L0_GI** | Sm$^R$, pUC, Lv0 GGtλT0-L0 terminator, GI fusion sites | This work |
| **pRK106-rnpB_T1_GI** | Sm$^R$, pUC, Lv0 rnpB_T1 terminator, GI fusion sites | This work |
| **pRK108-T500_GI** | Sm$^R$, pUC, Lv0 T500 terminator, GI fusion sites | This work |
| **pSS14-λT1_GI** | Sm$^R$, pUC, Lv0 λT1 terminator, GI fusion sites | This work |
| **pSEVA2219g2-pem7stGFP** | Km$^R$, RK2, Lv1 pSEVA2219g2 vector containing Pem7 promoter, RBS standard, *gfp* gene and BBa_B1006 terminator | This work |
| **pSEVA23g19g2-pem7stGFP** | Km$^R$, pBBR1, Lv1 pSEVA23g19g2 vector containing Pem7 promoter, RBS standard, *gfp* gene and BBa_B1006 terminator | This work |

| | | |
|---|---|---|
| **pSEVA2819g2-pem7stGFP** | Km$^R$, pUC, Lv1 pSEVA28-2 vector containing Pem7 promoter, RBS standard, *gfp* gene and BBa_B1006 terminator | This work |
| **pICH41295** | Sp$^R$/Sm$^R$, ColE1, MoClo L0 acceptor position AD with lacZα | (1) |
| **pICH41308** | Sp$^R$/Sm$^R$, ColE1, MoClo L0 acceptor position DG with lacZα | (1) |
| **pICH41276** | Sp$^R$/Sm$^R$, ColE1, MoClo L0 acceptor position GI with lacZα | (1) |
| **pL1F-1** | Amp$^R$, ColE1/RK2, Level 1 acceptor vector at position 1 with *lacZα* | (1) |
| **pL1F-2** | Amp$^R$, ColE1/RK2, Level 1 acceptor vector at position 2 with *lacZα* | (1) |
| **pL1F-3** | Amp$^R$, ColE1/RK2, Level 1 acceptor vector at position 3 with *lacZα* | (1) |
| **pL1F-1 *phaC1*** | Amp$^R$, ColE1/RK2, Lv1 pL1F-1 containing λT0-SynPro16, phaC1 and rnpB_T1 | This work |
| **pL1F-2 *phaA*** | Amp$^R$, ColE1/RK2, Lv1 pL1F-2 containing λT0-SynPro16, phaA and rpoC | This work |
| **pL1F-3 *phaB1*** | Amp$^R$, ColE1/RK2, Lv1 pL1F-3 containing λT0-SynPro16, phaB1 and T500 | This work |
| **pSEVA63g19gB PHB** | Gm$^R$, pBBR1, Lv2 pSEVA63g19gB vector containing pL1F-1 *phaC1*, pL1F-2 *phaA* and pL1F-3 *phaB1*. | This work |
| **pSEVA6219gB PHB** | Gm$^R$, RK2, Lv2 pSEVA6219gB vector containing pL1F-1 *phaC1*, pL1F-2 *phaA* and pL1F-3 *phaB1*. | This work |
| **pSEVA6819gB PHB** | Gm$^R$, pUC, Lv2 pSEVA6819gB vector containing pL1F-1 *phaC1*, pL1F-2 *phaA* and pL1F-3 *phaB1*. | This work |
| **Lv1 crtE** | Km$^R$, pBBR1, Lv1 pSEVA23g19g1 vector containing mix of promoters (J23100, J23106, J23107, J23116), RBS standard, *crtE* gene and GGtλT0-L0 terminator | This work |
| **Lv1 crtB** | Km$^R$, pBBR1, Lv1 pSEVA23g19g2 vector containing mix of promoters (J23100, J23106, J23107, J23116), RBS standard, *crtB* gene and rnpB_T1 terminator | This work |
| **Lv1 crtI** | Km$^R$, pBBR1, Lv1 pSEVA23g19g3 vector containing J23103 promoter, RBS standard, *crtB* gene and rnpB_T1 terminator | This work |
| **Lv1 crtY** | Km$^R$, pBBR1, Lv1 pSEVA23g19g4 vector containing mix of promoters (J23100, J23106, J23107, J23116), RBS standard, *crtB* gene and T500 terminator | This work |
| **Lv1 crtZ** | Km$^R$, pBBR1, Lv1 pSEVA23g19g5 vector containing mix of promoters (J23100, J23106, J23107, J23116), RBS standard, *crtB* gene and λT1 terminator | This work |
| **Lv2 zeaxanthin pUC** | Gm$^R$, pUC, Lv2 pSEVA6819gD vector containing mix of Lv1 crtE, Lv1 crtB, Lv1 crtI, Lv1 crtY and Lv1 crtZ | This work |
| **Lv2 zeaxanthin pBBR1** | Gm$^R$, pBBR1, Lv2 pSEVA63g19gD vector containing mix of Lv1 crtE, Lv1 crtB, Lv1 crtI, Lv1 crtY and Lv1 crtZ | This work |
| **Lv1 OleD** | Km$^R$, pBBR1, Lv1 pSEVA23g19g1 vector containing Trc promoter with LacO, RBS T7 from pET28, *OleD* gene and T7 terminator from pET28 | This work |
| **Lv1 OleD-C-GFP** | Km$^R$, pBBR1, Lv1 pSEVA23g19g1 vector containing Trc promoter with LacO, RBS T7 from pET28, *OleD* gene fussion with C-terminal GFP and T7 terminator from pET28 | This work |

| | | |
|---|---|---|
| **Lv1 N-His-OleD-C-GFP** | Km$^R$, pBBR1, Lv1 pSEVA23g19g1 vector containing Trc promoter with LacO, RBS T7 from pET28, *OleD* gene fussion with N-terminal 6xHis tag and C-terminal GFP, and T7 terminator from pET28 | This work |
| **Lv1 N-Strep-OleD-C-GFP** | Km$^R$, pBBR1, Lv1 pSEVA23g19g1 vector containing Trc promoter with LacO, RBS T7 from pET28, *OleD* gene fussion with N-terminal Strep tag and C-terminal GFP, and T7 terminator from pET28 | This work |
| **Lv1 N-GST-OleD-C-GFP** | Km$^R$, pBBR1, Lv1 pSEVA23g19g1 vector containing Trc promoter with LacO, RBS T7 from pET28, *OleD* gene fussion with N-terminal GST tag and C-terminal GFP, and T7 terminator from pET28 | This work |
| **Lv1 N-MBP-OleD-C-GFP** | Km$^R$, pBBR1, Lv1 pSEVA23g19g1 vector containing Trc promoter with LacO, RBS T7 from pET28, *OleD* gene fussion with N-terminal MBP tag and C-terminal GFP, and T7 terminator from pET28 | This work |

Table S4. List of oligonucleotides used in this study

| Primer | Sequence (5'-3') | Comments |
|---|---|---|
| **PS1** | AGGGCGGCGGATTTGTCC | Sequencing GS plasmids |
| **PS2** | GCGGCAACCGAGCGTTC | Sequencing GS plasmids |
| **R24** | AGCGGATAACAATTTCACACAGGA | Sequencing GS plasmids |
| **F24** | CGCCAGGGTTTTCCCAGTCACGAC | Sequencing GS plasmids |
| **5pBBR1_1** | CTCTGCGAGGCTGGCCGTAGG | Removing BpI site |
| **3pBBR1_1** | GCCGTGGTGGTCAGCCAGAAAACACTTTCCAAGCTCATCGG | Removing BpI site |
| **5pBBR1_2** | CCGATGAGCTTGGAAAGTGTTTTCTGGCTGACCACCACGGC | Removing BpI site |
| **3pBBR1_2** | CCTAGACAGCTGGGCGCGCC | Removing BpI site |
| **primer 1A** | GGGAAGACAATGCCGGAGTGAGACCGCAGCTGGCACGACAGGTTTGC | Construction Lv1 |
| **primer I2** | GAAGACAATTGCAGCGTGAGACCGTC | Construction Lv1 |
| **primer 2A** | GGGAAGACAAGCAAGGAGTGAGACCGCAGCTGGCACGACAGGTTTGC | Construction Lv1 |
| **primer I3** | GAAGACAATAGTAGCGTGAGACCGTCACAG | Construction Lv1 |
| **primer 3A** | GGGAAGACAAACTAGGAGTGAGACCGCAGCTGGCACGACAGGTTTGC | Construction Lv1 |
| **primer I4** | GAAGACAAGTAAAGCGTGAGACCGTCACAG | Construction Lv1 |
| **primer 4A** | GGGAAGACAATTACGGAGTGAGACCGCAGCTGGCACGACAGGTTTGC | Construction Lv1 |
| **primer I5** | GAAGACAATCTGAGCGTGAGACCGTCAC | Construction Lv1 |
| **primer 5A** | GGGAAGACAACAGAGGAGTGAGACCGCAGCTGGCACGACAGGTTTGC | Construction Lv1 |
| **primer I6** | GAAGACAACACAAGCGTGAGACCGTCAC | Construction Lv1 |

| | | |
|---|---|---|
| **primer 6A** | GGGAAGACAATGTGGGAGTGAGACCGCAGCTGGC ACGACAGGTTTGC | Construction Lv1 |
| **primer I7** | GAAGACAAGCTCAGCGTGAGACCGTC | Construction Lv1 |
| **primer 1I** | GGGAAGACAATGCCAGCGTGAGACCGCAGCAGGC ACGACAGGTTTGC | Construction Lv1 |
| **primer A2** | GAAGACAATTGCGGAGTGAGACCGTCACAGCTTGT C | Construction Lv1 |
| **A1** | GGGTCTCAGGAGTGCCTTGTCTTCACGACAGGTTT GCCGACTGG | Construction Lv2 |
| **B1** | GGGTCTCAGGAGTGCCTTGTCTTCACGACAGGTTT GCCGACTGG | Construction Lv2 |
| **C1** | GGGTCTCACCATTGCCTTGTCTTCACGACAGGTTT GCCGACTGG | Construction Lv2 |
| **D1** | GGGTCTCAAATGTGCCTTGTCTTCACGACAGGTTT GCCGACTGG | Construction Lv2 |
| **E1** | GGGTCTCAAGGTTGCCTTGTCTTCACGACAGGTTT GCCGACTGG | Construction Lv2 |
| **3B** | GGTCTCAAGTATAGTTTGTCTTCTCACAGCTTGTCT GTAAGCGGATGC | Construction Lv2 |
| **4C** | GGTCTCAATGGGTAATTGTCTTCTCACAGCTTGTCT GTAAGCGGATGC | Construction Lv2 |
| **5D** | GGTCTCACATTTCTGTTGTCTTCTCACAGCTTGTCT GTAAGCGGATGC | Construction Lv2 |
| **6E** | GGTCTCAACCTCACATTGTCTTCTCACAGCTTGTCT GTAAGCGGATGC | Construction Lv2 |
| **7F** | GGTCTCACGAAGCTCTTGTCTTCTCACAGCTTGTCT GTAAGCGGATGC | Construction Lv2 |
| **C2** | CCGAAGACAATTGCATGGTGAGACCGTCACAGCTT GTCTGTAAGCGGTG | Construction Lv3 |
| **D3** | CCGAAGACAATAGTCATTTGAGACCGTCACAGCTT GTCTGTAAGCGG | Construction Lv3 |
| **E4** | CCGAAGACAAGTAAACCTTGAGACCGTCACAGCTT GTCTGTAAGCGG | Construction Lv3 |
| **F5** | CCGAAGACAATCTGCGAATGAGACCGTCACAGCTT GTCTGTAAGCGG | Construction Lv3 |
| **RK81** | GGAAGAGCGCCCAATACG | Sequencing of level 0 MoClo plasmids |
| **RK82** | AAAGTGCCACCTGACGTCTA | Sequencing of level 0 MoClo plasmids |
| **RK155** | GAACCCTGTGGTTGGCATGCACATAC | Sequencing of level 1 MoClo plasmids |
| **RK156** | CTGGTGGCAGGATATATTGTGGTG | Sequencing of level 1 MoClo plasmids |
| **RK246** | TTTGAAGACACGGAGCTTGGACTCCTGTTGATAGA TCC | λT0-SynPro16 Level 0 MoClo part |
| **RK247** | TTTGAAGACACCATTTAGAACCCCCTCGTACGCTC | λT0-SynPro16 Level 0 MoClo part |
| **SS15** | TTTGAAGACGAAATGGCGACCGGCAAAGGCGCGG CA | *C. necator phaC1 H16_A1437* Level 0 MoClo part |

| | | |
|---|---|---|
| **SS16** | TTTGAAGACGAGTCCTCCATCATGTTGCGCACGCCGGC | *C. necator phaC1 H16_A1437* Level 0 MoClo part |
| **SS17** | TTTGAAGACAGGGACCTGACACGCGGCAAGATCT | *C. necator phaC1 H16_A1437* Level 0 MoClo part |
| **SS18** | TTTGAAGACAGAACACCACGGCGCCTTCGGTCAC | *C. necator phaC1 H16_A1437* Level 0 MoClo part |
| **SS19** | TTTGAAGACGTTGTTCGAGAACGAGTACTTCCAGCT | *C. necator phaC1 H16_A1437* Level 0 MoClo part |
| **SS20** | TTTGAAGACGTCCTCGCGCGAGCCGTAGATATA | *C. necator phaC1 H16_A1437* Level 0 MoClo part |
| **SS21** | TTTGAAGACGAGAGGACCATATCGTGCCGTGGACC | *C. necator phaC1 H16_A1437* Level 0 MoClo part |
| **SS22** | TTTGAAGACGAAAGCTCATGCCTTGGCTTTGACGTATCG | *C. necator phaC1 H16_A1437* Level 0 MoClo part |
| **SS9** | TTTGAAGACGAAATGACTGACGTTGTCATCGTATCCGC | *C. necator phaA H16_A1438* Level 0 MoClo part |
| **SS10** | TTTGAAGACGAAAGCTTATTTGCGCTCGACTGCCAG | *C. necator phaA H16_A1438* Level 0 MoClo part |
| **SS11** | TTTGAAGACCTAATGACTCAGCGCATTGCGTATGTGACCG | *C. necator phaB1 H16_A1439* Level 0 MoClo part |
| **SS12** | TTTGAAGACCTGACACCGTGTTGACGGTCACGCC | *C. necator phaB1 H16_A1439* Level 0 MoClo part |
| **SS13** | TTTGAAGACGATGTCTCCGGGCTATATCGCCACCGACA | *C. necator phaB1 H16_A1439* Level 0 MoClo part |
| **SS14** | TTTGAAGACGAAAGCTCAGCCCATATGCAGGCCGCC | *C. necator phaB1 H16_A1439* Level 0 MoClo part |
| **RK69** | TTTGAAGACCTGCTTCTTGGACTCCTGTTGATAGA | λT0 Level 0 MoClo part |
| **RK70** | TTTGAAGACCTAGCGCTGGATTCTCACCAATAAAAAAC | λT0 Level 0 MoClo part |
| **RK189** | TTTGAAGACGAGCTTTCTAGGGCGGCGGATTTGTC | T1 (*rrnB)* Level 0 MoClo part |
| **RK190** | TTTGAAGACGAAGCGGGCATCAAATAAAACGAAAGGC | T1 (*rrnB)* Level 0 MoClo part |
| **RK205** | TTTGAAGACAGGCTTTCGGTCAGTTTCACCTGATT | *rnpB*_T1 Level 0 MoClo part |
| **RK206** | TTTGAAGACAGAGCGGACAGTCATTCATCTTTCTGCC | *rnpB*_T1 Level 0 MoClo part |
| **RK207** | TTTGAAGACGTGCTTGTAATCGTTAATCCGCAAATAA | *rpoC* Level 0 MoClo part |

| | | |
|---|---|---|
| **RK208** | TTTGAAGACGTAGCGTGACAAATGCTCTTTCCCTA | *rpoC* Level 0 MoClo part |
| **RK209** | TTTGAAGACTCGCTTCAAAGCCCGCCGAAAGG | T500 Level 0 MoClo part |
| **RK210** | TTTGAAGACTCAGCGACAGAAAAGCCCGCCTTTCG | T500 Level 0 MoClo part |
| **MM319** | GAACGACCTGGTGTGGAACTAC | Sequencing of level 2 PHB- GS plasmids |
| **MM320** | CAACTTCCTTGCCACCAATCC | Sequencing of level 2 PHB- GS plasmids |
| **MM325** | CAGCATGTCCGGCCTCAAG | Sequencing of level 2 PHB- GS plasmids |
| **MM346** | GACTCCTCCGACGACAACC | Sequencing of level 2 PHB- GS plasmids |
| **SS9** | TTTGAAGACGAAATGACTGACGTTGTCATCGTATCC GC | Sequencing of level 2 PHB- GS plasmids |
| **SS10** | TTTGAAGACGAAAGCTTATTTGCGCTCGACTGCCA G | Sequencing of level 2 PHB- GS plasmids |

**Table S5.** GC-MS analysis of PHB content at 24 h in PHB production conditions. Mean values and standard deviations of three biological replicates with two technical replicates for each. CDW: cell dry weight. See Table S2 for plasmid descriptions.

| Strain (*ori*V) | 24 h $OD_{600}$ | Total CDW (g $L^{-1}$) | PHB (% CDW) |
|---|---|---|---|
| ***E. coli*** **DH10B [pSEVA6819gB PHB] (pUC)** | 21.60 ± 0.2 | 5.37 ± 0.07 | 50.23 ± 2.28 |
| ***E. coli*** **DH10B [pSEVA63g19gB PHB] (pBBR1)** | 6.37 ± 0.15 | 1.63 ± 0.02 | 12.90 ± 0.74 |
| ***E. coli*** **DH10B [pSEVA6219gB PHB] (RK2)** | 5.57 ± 0.06 | 1.47 ± 0.08 | 7.67 ± 0.99 |

e16765.

2. Werner,S., Engler,C., Weber,E., Gruetzner,R. and Marillonnet,S. (2012) Fast track assembly of multigene constructs using golden gate cloning and the MoClo system. *Bioeng. Bugs*, **3**, 38–43.
3. Tiso,T., Sabelhaus,P., Behrens,B., Wittgens,A., Rosenau,F., Hayen,H. and Blank,L.M. (2016) Creating metabolic demand as an engineering strategy in Pseudomonas putida – Rhamnolipid synthesis as an example. *Metab. Eng. Commun.*, **3**, 234–244.
4. Martínez-García,E., Aparicio,T., de Lorenzo,V. and Nikel,P.I. (2014) New transposon tools tailored for metabolic engineering of Gram-negative microbial cell factories. *Front. Bioeng. Biotechnol.*, **2**.
5. Revelles,O., Tarazona,N., García,J.L. and Prieto,M.A. (2016) Carbon roadmap from syngas to polyhydroxyalkanoates in Rhodospirillum rubrum. *Environ. Microbiol.*, **18**, 708–720.
6. Tronina,T., Bartmańska,A., Milczarek,M., Wietrzyk,J., Popłoński,J., Rój,E. and Huszcza,E. (2013) Antioxidant and antiproliferative activity of glycosides obtained by biotransformation of xanthohumol. *Bioorganic Med. Chem. Lett.*, **23**, 1957–1960.
7. Zobel,S., Benedetti,I., Eisenbach,L., De Lorenzo,V., Wierckx,N. and Blank,L.M. (2015) Tn7-Based Device for Calibrated Heterologous Gene Expression in Pseudomonas putida. *ACS Synth. Biol.*, **4**, 1341–1351.
8. Torres-Bacete,J., Luís García,J. and Nogales,J. (2021) A portable library of phosphate-depletion based synthetic promoters for customable and automata control of gene expression in bacteria. *Microb. Biotechnol.*, 10.1111/1751-7915.13808.
9. Nogales,J., Canales,Á., Jiménez-Barbero,J., Serra,B., Pingarrón,J.M., García,J.L. and Díaz,E. (2011) Unravelling the gallic acid degradation pathway in bacteria: The gal cluster from Pseudomonas putida. *Mol. Microbiol.*, **79**, 359–374.
10. Durfee,T., Nelson,R., Baldwin,S., Plunkett,G., Burland,V., Mau,B., Petrosino,J.F., Qin,X., Muzny,D.M., Ayele,M., *et al.* (2008) The complete genome sequence of Escherichia coli DH10B: Insights into the biology of a laboratory workhorse. *J. Bacteriol.*, **190**, 2597–2606.
11. Regenhardt,D., Heuer,H., Heim,S., Fernandez,D.U., Strömpl,C., Moore,E.R.B. and Timmis,K.N. (2002) Pedigree and taxonomic credentials of Pseudomonas

putida strain KT2440. *Environ. Microbiol.*, **4**, 912–915.
12. Iverson,S. V., Haddock,T.L., Beal,J. and Densmore,D.M. (2016) CIDAR MoClo: Improved MoClo Assembly Standard and New E. coli Part Library Enable Rapid Combinatorial Design for Synthetic and Traditional Biology. *ACS Synth. Biol.*, **5**, 99–103.
